# Catecholamines Alter the Intrinsic Variability of Cortical Population Activity and Perception

**DOI:** 10.1101/170613

**Authors:** Thomas Pfeffer, Arthur-Ervin Avramiea, Guido Nolte, Andreas K. Engel, Klaus Linkenkaer-Hansen, Tobias H. Donner

## Abstract

The ascending modulatory systems of the brainstem are powerful regulators of global brain state. Disturbances of these systems are implicated in several major neuropsychiatric disorders. Yet, how these systems interact with specific neural computations in the cerebral cortex to shape perception, cognition, and behavior remains poorly understood. Here, we probed into the effect of two such systems, the catecholaminergic (dopaminergic and noradrenergic) and cholinergic systems, on an important aspect of cortical computation: its intrinsic variability. To this end, we combined placebo-controlled pharmacological intervention in humans, magnetoencephalographic (MEG) recordings of cortical population activity, and psychophysical measurements of the perception of ambiguous visual input. A low-dose catecholaminergic, but not cholinergic, manipulation altered the rate of spontaneous perceptual fluctuations as well as the temporal structure of “scale-free” population activity of large swaths of visual and parietal cortex. Computational analyses indicate that both effects were consistent with an increase in excitatory relative to inhibitory activity in the cortical areas underlying visual perceptual inference. We propose that catecholamines regulate the variability of perception and cognition through dynamically changing the cortical excitation-inhibition ratio. The combined read-out of fluctuations in perception and cortical activity we established here may prove useful as an efficient, and easily accessible marker of altered cortical computation in neuropsychiatric disorders.

## INTRODUCTION

The modulatory systems of the brainstem send widespread, ascending projections to the specialized circuits of the cerebral cortex that mediate perception, cognition, and goal-directed behavior. These systems regulate ongoing changes in brain state, even during periods of wakefulness [1–4]. They are recruited during, and in turn shape, cognitive processes, such as perceptual inference, learning, and decision-making [5–8]. Because these systems are implicated in most neuropsychiatric disorders, they are also major targets of pharmacological therapy of brain disorders [5,9,10]. Taken together, neuromodulatory systems have remarkably specific effects on cognition, despite the widespread and diffuse nature of their projections to cortex. An important challenge for neuroscience is to uncover the mechanistic principles, by which neuromodulatory systems interact with the cortical computations underlying cognition.

One key parameter of cortical computation that might be under neuromodulatory control is the intrinsic variability – i.e., fluctuations that occur during constant (or absent) sensory input [13,14]. Specifically, it has been proposed that the catecholaminergic neuromodulators noradrenaline and dopamine may shift the cortical computations underlying decision-making from a stable (“exploitative”) to a variable (“exploratory”) mode [5,15]. A context-dependent adjustment of the variability of cortical computations may also be adaptive for perceptual inference in the face of ambiguous sensory input [16].

Animal work has shown that catecholamines and acetylcholine, another important neuromodulator, alter the intrinsic variability of neural activity [2,11,17–19] through highly selective interactions with specific elements (pyramidal cells and/or inhibitory interneurons) of cortical microcircuits [20,21]. But it is unknown how these changes at the level of cortical microcircuits relate to the intrinsic variability of perception and cognition.

At the larger scale of cortical mass action that is assessable with non-invasive recordings in humans, activity also fluctuates intrinsically, in a spatially and temporally structured manner [22,23]. The temporal structure of these fluctuations is characteristic of so-called “scale-free” behavior: Power spectra that scale as a function of frequency according to a power law, P(f) ∝ f^*β*^ [24,25], indicating long-range temporal autocorrelations [26–29]. Some studies have linked the spatio-temporal structure of the fluctuations in cortical population activity to specific perceptual and cognitive processes [28,30–32]. But it is unknown if and how these fluctuations in cortical population activity are dynamically regulated by neuromodulatory systems.

We aimed to close these gaps by systematically quantifying the effects of catecholaminergic and cholinergic neuromodulation on the intrinsic variability in perception and large-scale cortical activity in the healthy human brain. To this end, we combined placebo-controlled, selective pharmacological interventions, psychophysical measurements of fluctuations in perception in the face of a continuously presented and ambiguous visual stimulus, and MEG recordings of fluctuations in cortical population activity.

Catecholamines, but not acetylcholine, increased both the variability of perception as well as long-range temporal correlations of intrinsic cortical activity in visual and parietal cortex. Based on previous theoretical and experimental work [33–36], we interpreted the increase in perceptual variability in terms of an increase in the net ratio between cortical excitation and inhibition in those cortical regions. Simulating a recurrent neural network under synaptic gain modulation enabled us to show that an analogous mechanism may account for the increase of long-range temporal correlations of cortical activity under catecholamines.

## RESULTS

We tested for changes in intrinsic fluctuations of perception and cortical population activity under placebo-controlled, within-subjects pharmacological manipulations of catecholamine (using the noradrenaline reuptake inhibitor atomoxetine) and acetylcholine (using the cholinesterase inhibitor donepezil) levels (Fig 1A, see Methods for details on the pharmacological interventions).

**Fig 1.**
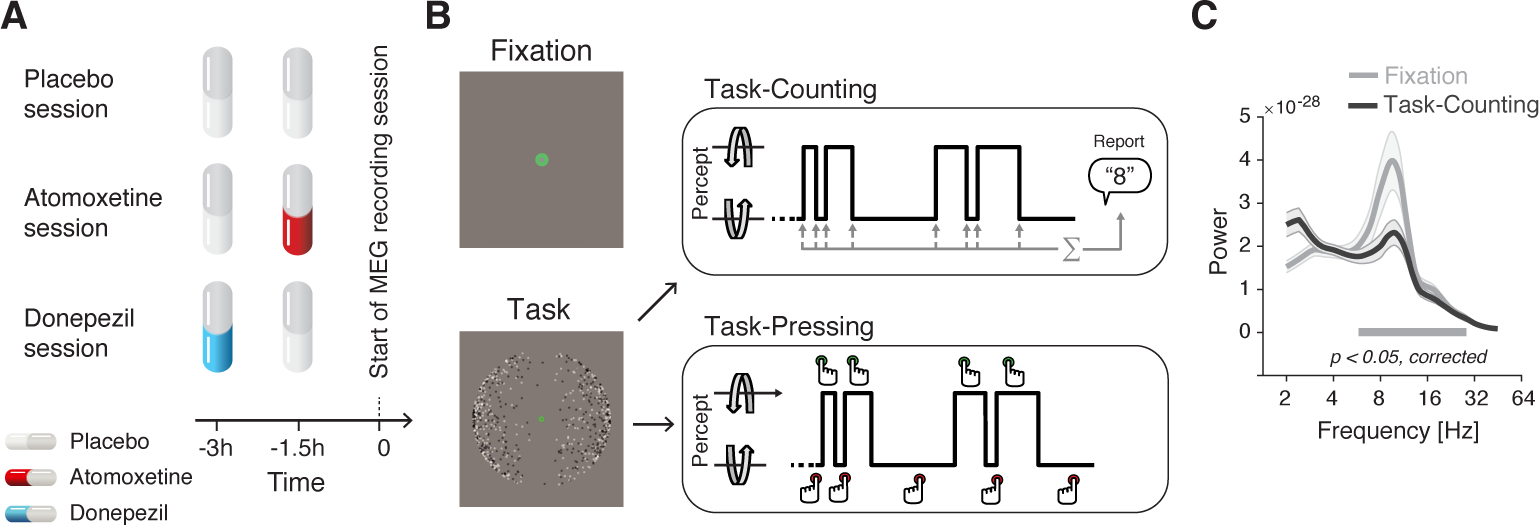
Experimental design **(A, B)** Types and time course of experimental sessions. **(A)** Each subject participated in three sessions, involving administration of placebo, atomoxetine, or donepezil (session order randomized across subjects). Each session entailed the administration of two pills, in the order depicted for the different session types. **(B)** Within each session, subjects alternated between three conditions, Fixation, Task-Counting and Task-Pressing, during which MEG was recorded (runs of 10 min each). See Materials and Methods for details. **(C)** Group average power spectrum, averaged across all MEG sensors, for Rest and Task (Placebo condition only).

Fluctuations in cortical activity were measured during two steady-state conditions, both of which excluded transients in sensory input or motor output (Fig 1B): (i) fixation of an otherwise gray screen (a condition termed “Fixation”), as in most studies of human “resting-state” activity [22,23]; and (ii) silent counting of the spontaneous alternations in the perceptual interpretation of a continuously presented, ambiguous visual stimulus (dubbed “Task-counting”). In a third condition that was only used for the analysis of perceptual fluctuations, subjects immediately reported the perceptual alternations by button press (“Task-pressing”; i.e. associated with movement-related transients in cortical activity). This design capitalized on recent insights into the circuit mechanisms underlying intrinsic perceptual dynamics [33,34,36] which helped constrain the mechanistic interpretation of the results reported below.

The Results section is organized as follows. We first present the effects of the “Atomoxetine” and “Donepezil” conditions (each compared against the “Placebo” condition) on the rate of perceptual fluctuations. These effects were in line with a boost in the relative strength of excitatory drive of visual cortex under Atomoxetine. We then show how (i) constant sensory and task drive (i.e., Task-counting vs. Fixation) and (ii) the pharmacological manipulations affect the intrinsic fluctuations in cortical activity. We focus on the temporal auto-correlation structure of intrinsic fluctuations in the amplitude of band-limited cortical population activity (see Methods and Fig 2). Control analyses showing the drug effects on other measures of cortical population activity and peripheral physiological signals support the validity and specificity of our conclusions.

**Fig 2.**
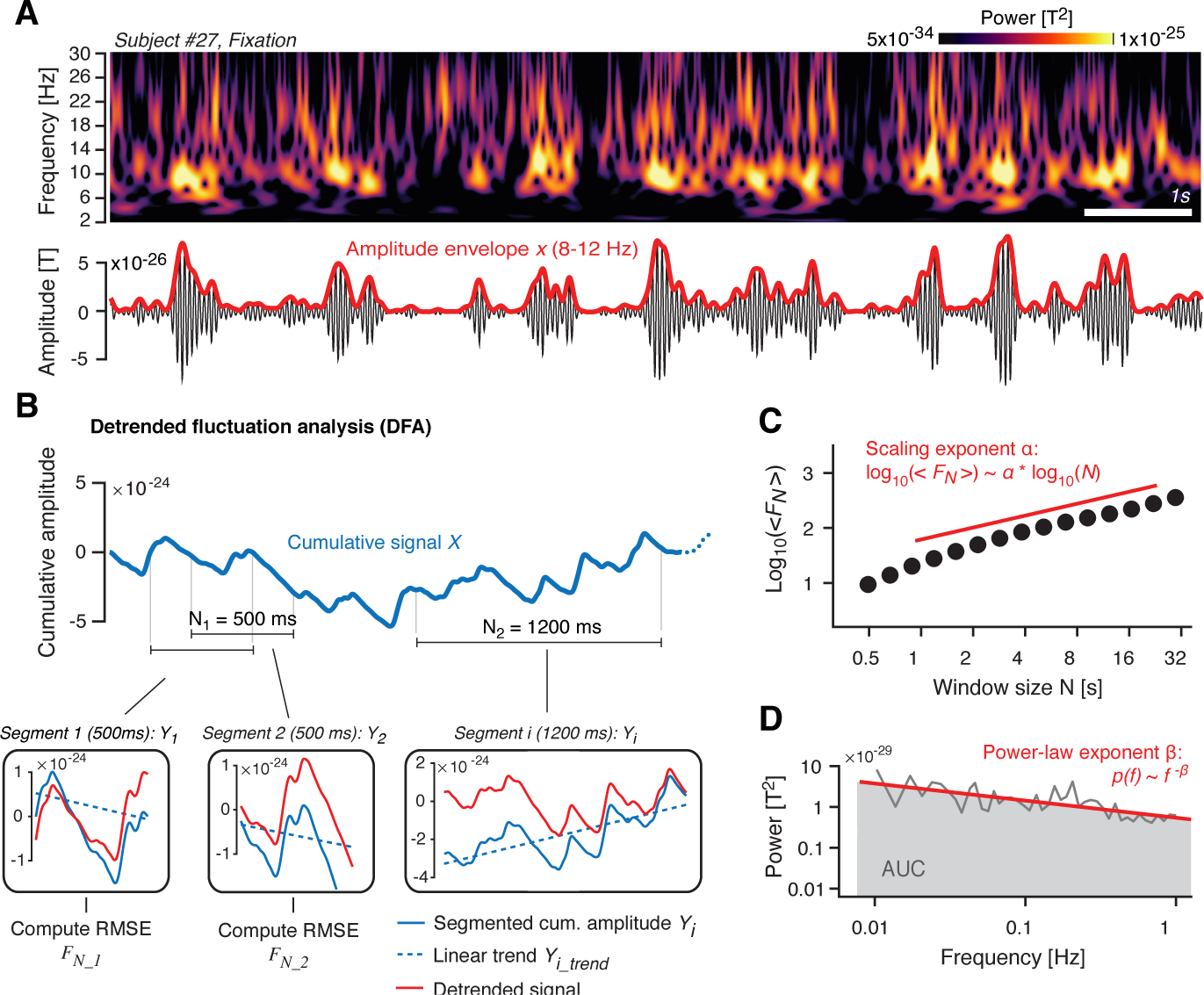
Quantifying the temporal structure of fluctuations in oscillatory cortical activity. **(A)** *Top*. Time-frequency representation of MEG power fluctuations during Rest (example subject). *Bottom*. Filtered signal (10 Hz; black) and the corresponding amplitude envelope (red). **(B)** Illustration of detrended fluctuation analysis. See main text (Materials and Methods) for details. *Top*. Cumulative sum of the amplitude envelope. *Bottom*. Detrending of cumulative sum within segments, shown for two different window lengths *N* (N_1_ = 500 ms and N_2_ = 1200 ms). **(C)** Root-mean-square fluctuation function <*F_N_*>. In log-log coordinates, <*F_N_*> increases approximately linearly as a function of *N*, with a slope that is the scaling exponent *α*. **(D)** Illustration of power spectrum analysis of amplitude envelope. In log-log coordinates, the power spectrum can be approximated by a straight line, with a slope *β* (power-law exponent) and an area under the curve (gray) that quantifies the overall variance of the signal.

We close with simulations of a patch of recurrently connected excitatory and inhibitory integrate-and-fire neurons. The simulations show that the changes in temporal correlations observed in the MEG data can be explained by a modulation synaptic gain that altered the net ratio between excitatory and inhibitory activity.

### Atomoxetine increases the rate of bistable perceptual fluctuations

The ambiguous visual stimulus that was continuously presented during both Task-counting and Task-pressing induced ongoing fluctuations in perception, i.e., spontaneous alternations between two apparent rotation directions of 3D-motion (Fig 1B; see Movie M1), a phenomenon is referred to as multi-stable perception. The rate of the perceptual alternations reported by the participants provided a read-out of visual cortical circuit state. Current models explain bistable perceptual fluctuations in terms of the interplay between feedforward, excitatory drive of stimulus-selective neural populations in visual cortex, mutual inhibition between them, stimulus-selective adaptation, and neural “noise” [33,34]. Increases in the ratio between feedforward, excitatory input to, and mutual inhibition within the cortical circuit, give rise to faster perceptual alternations. This idea is supported by convergent evidence from functional magnetic resonance imaging, magnetic resonance spectroscopy, and pharmacological manipulation of GABAergic transmission [31,36]. We reasoned that neuromodulators such as noradrenaline might dynamically change these parameters [12,37], and thereby alter the rate of perceptual fluctuations.

Atomoxetine increased the rate of perceptual fluctuations compared to both Placebo and Donepezil conditions (Fig 3A; Atomoxetine vs. Placebo: *p* = 0.007; *t* = 2.913; Atomoxetine vs. Donepezil: *p* = 0.001; *t* = 3.632; Donepezil vs. Placebo: *p* = 0.966; *t* = −0.043; all paired t-tests, pooled across Task-counting and Task-pressing). The perceptual alternation rates were highly consistent across Task-counting and Task-pressing (Fig S1C), supporting the validity of the counting condition as behavioral read-out of bistable perceptual fluctuations. Likewise, the Atomoxetine effect on perceptual fluctuation rate was evident for Task-counting (*p* = 0.045; *t* = 2.103; paired t-test; Fig S1A) and Task-pressing (*p* = 0.018; *t* = 2.540; paired t-test; Fig S1B) individually.

**Fig 3.**
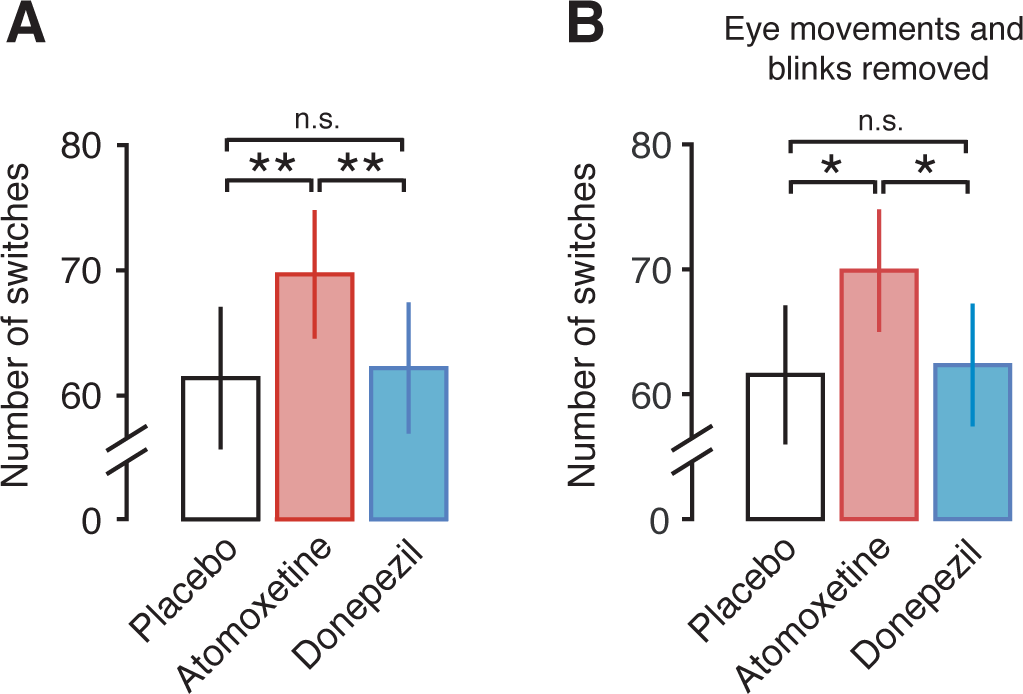
Atomoxetine, but not Donepezil, increases the rate of perceptual alternations **(A)** Number of perceptual alternations reported by the subjects per 10 min run, pooled across task conditions (Task-counting and Task-pressing). **(B)** Same as (A), after regressing out blink and eye movement data (see Methods and Supplementary Figure S2). Significance was assessed using two-sided paired t-tests (N=28).

These changes in perceptual fluctuations were not explained by an increase in the rates of eye blinks or fixational eye movements. First, there was no significant increase during Atomoxetine compared to Placebo in any of five different eye movement parameters measured here (Fig S2). Second, none of these parameters correlated with the perceptual alternation rate (Fig S2). Third, and most importantly, the effect of Atomoxetine on the perceptual dynamics was also significant after removing (via linear regression) the individual eye movement parameters (Fig 3B).

In sum, Atomoxetine had an effect on bistable perceptual fluctuations that was both robust and specific, evident when compared with either Placebo or Donepezil. This effect was in line with an increase in the strength of excitatory feedforward drive of visual cortex relative to the strength of mutual inhibition between the neural sub-populations encoding the competing perceptual interpretations of the ambiguous stimulus. Such an effect should have occurred in the motion-sensitive visual cortical areas, which implement the visual competition induced by the ambiguous structure-from-motion stimulus [38,39].

### Atomoxetine increases the scaling exponent of fluctuations in cortical population activity

We estimated long-range temporal correlations of band-limited amplitude fluctuations (indicated by the scaling exponent *α*; see Methods for details) to quantify intrinsic fluctuations in cortical population activity. Our analyses focused on amplitude envelope fluctuations in the 8–12 Hz frequency range (“alpha band”), for two reasons. First, as expected from previous work [40], the cortical power spectra exhibited a clearly discernible peak in this frequency range, which robustly modulated with sensory or task drive (suppressed under Task-counting, Fig 1C). Second, previous studies reported robust long-range temporal correlations with peaks in the same frequency range [26,29].

We first replicated two previously established observations pertaining to the scaling exponent *α*. First, the average across cortical patches and participants was *α* = 0.67 (s.d. = ±0.09) during Fixation (Placebo only) and *α* = 0.64 (s.d. = ±0.07) during Task-counting (Placebo only), indicative of long-range temporal correlations similar to the ones found in previous work [26,29,41]. Second, the sensory and task drive during Task-counting reliably reduced *α* compared to Fixation, again as shown in previous work [27,42]. Across all voxels, *α* was significantly larger during Fixation than during Task-counting (*p* = 0.0062; *t* = 2.97; paired t-test; Placebo condition only). This difference was significant across pharmacological conditions in large parts of cortex including the occipital and parietal regions that were driven by the motion stimulus (*p* < 0.05; cluster-based permutation test; Fig 4D).

**Fig 4.**
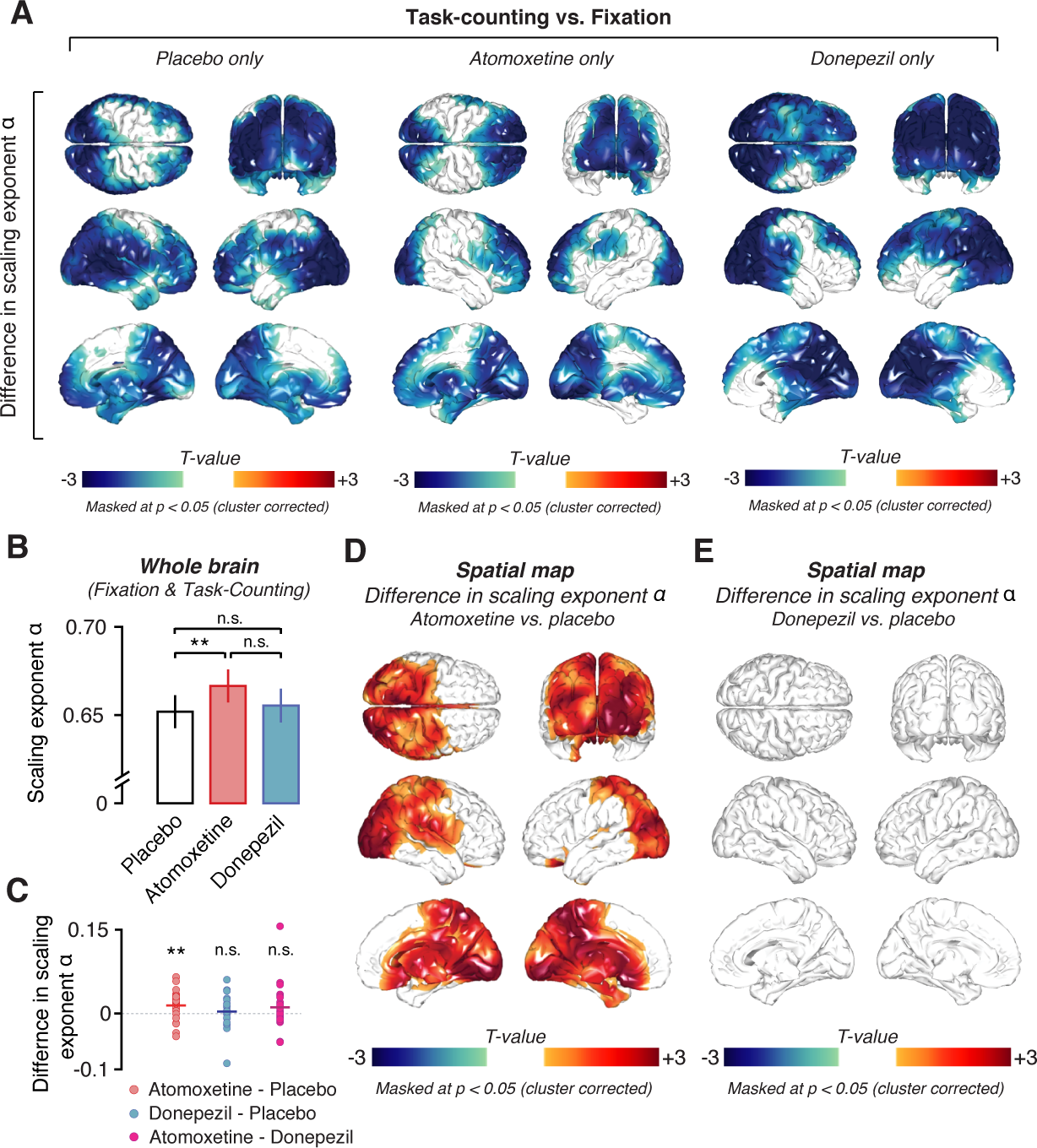
Effects of task and pharmacological conditions on long-range temporal correlations of the amplitude envelope of 8-12 Hz MEG activity. **(A)** Spatial distribution of significant differences in scaling exponent *α* between Task-counting and Fixation during Placebo (left), Atomoxetine (middle), and Donepezil (right). **(B)** Comparison between mean scaling exponents *α* averaged across all the entire brain (see Methods) during the different pharmacological conditions. **(C)** Individual subject differences in scaling exponent *α* between all drug conditions. **(D, E)** Spatial distribution of drug-induced changes in scaling exponents. **(D)** Atomoxetine vs. Placebo. **(E)** Donepezil vs. Placebo. Two-sided permutation tests (N=28); all statistical maps: Threshold at *p* = 0.05, cluster-based. All drug comparisons are averaged across behavioral conditions, i.e., Fixation and Task-counting.

Having verified the validity of our measurements of *α* we then tested for changes in *α* under the pharmacological conditions (Fig 4B-E and Fig 5). There was a highly significant increase in *α* for Atomoxetine compared to Placebo when collapsing across voxels as well as across Fixation and Task-counting (*p* = 0.0068; *t* = 2.93; paired t-test, Fig 4B-C). This effect was widespread, but not homogenous across cortex, comprising occipital and posterior parietal, as well as a number of midline regions including the thalamus (Fig 4D, *p* = 0.0022; cluster-based permutation test). Because it is unclear to which extent intrinsic activations from deep sources can be recovered using MEG, we focus our description and conclusions on the effects in cortical regions. Importantly, the Atomoxetine effect on *α* was also present at the level of MEG sensors (Fig S4), and hence did not depend on the source reconstruction method applied here (see Methods).

**Fig 5.**
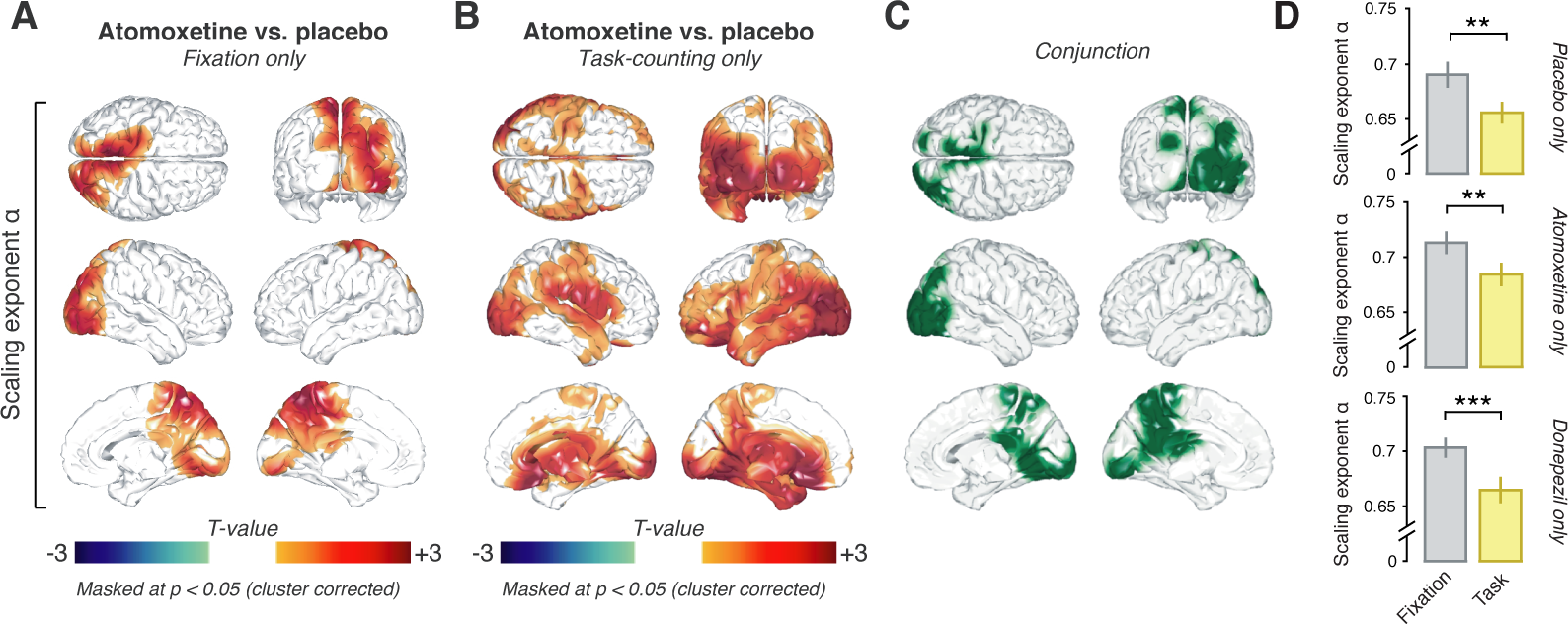
Atomoxetine increases long-range temporal correlations irrespective of behavioral condition. Spatial distribution of the Atomoxetine-induced changes in scaling exponent *α* during **(A)** Fixation and **(B)** Task-counting. **(C)** Conjunction of maps in (A) and (B), highlighting (in green) voxels with significant increases in both conditions. **(D)** Scaling exponents for Fixation (gray) and Task-counting (yellow) within conjunction cluster depicted in panel C for Placebo (top), Atomoxetine (middle) and Donepezil only (bottom).

The effect of Atomoxetine on *α* was subtle, likely due to the low dosage. But, importantly, the effect was highly reproducible across repeated measurements. We assessed reproducibility with two complementary approaches. The first was a region-of-interest (ROI) analysis. We defined a ROI in terms of a significant cluster for Atomoxetine > Placebo (one-sided paired t-test, *p* < 0.05, uncorrected) during the first run collected in each session (Fixation and Task-counting collapsed) and extracted this ROI’s mean *α* from the second run. We then reversed the procedure and so extracted a second, independent ROI-based *α* and averaged the *α*-estimates. This approach revealed a strong increase under Atomoxetine (*p* = 0.0023; *t* = 3.365). The second approach assessed the reproducibility of the spatial pattern of effects across both runs. To this end, we correlated the (non-thresholded) individual maps for the Atomoxetine vs. Placebo difference computed from the first and second run in each session (again pooling across Task-counting and Fixation) and tested the resulting correlation coefficients across participants. The average correlation was significantly different from zero (mean r = 0.29, p < 0.0001; permutation test against zero).

The Atomoxetine-related increases in scaling exponent *α* were evident during both Fixation and Task-counting (Fig 5A, Fixation: *p* = 0.0245; Fig 5B, Task-counting: *p* = 0.0035; cluster-based permutation test). The effects occurred in largely overlapping regions of occipital and parietal cortex (Fig 5C). There was no interaction between the effects of Atomoxetine and Task-counting anywhere in cortex: a direct comparison of the two Atomoxetine vs. Placebo difference maps, from Fixation and from Task-counting, yielded no significant clusters (*p* > 0.081 for all clusters; cluster-based permutation test). The same cortical regions in which *α* increased during Atomoxetine exhibited decreases during Task-counting: When testing for the task-dependent change in *α* (Fig 4A) specifically in the regions comprising the conjunction cluster of the Atomoxetine effect (Fig 5C), the reduction during Task-counting was also highly significant (Fig 5D) in all pharmacological conditions.

In contrast to the robust effect of Atomoxetine on *α*, there was no evidence for an effect of Donepezil at the dosage used here. The difference between Donepezil and Placebo (collapsed across Fixation and Task-counting) did not reach significance, neither when pooling across voxels (*p* = 0.50; *t* = 0.68; *BF* = 0.68; paired t-test; Fig 4B), nor when testing all voxels individually (*p* > 0.22 for all clusters; cluster-based permutation test; Fig 4E; Fig S5). Atomoxetine also increased the scaling exponents when directly compared to Donepezil during Task-counting (Fig S6A; *p* < 0.05; two-sided cluster-based permutation test), but not during Fixation (Fig S6B).

Taken together, the rich experimental design gave rise to a highly specific and consistent pattern of changes in *α* under the different experimental conditions, which helped constrain the mechanistic interpretation of the results. The Atomoxetine effects were specific, and not just due to the application of any drug targeting neurotransmitter systems. It is possible that the absence of detectable Donepezil effects on *α* was due to the low dosage or short administration period used here. However, the control analyses presented in the next section revealed clear effects of Donepezil on both cortical activity as well as markers of peripheral nervous system activity.

### Control analyses for the drug effects on other features of cortical dynamics or peripheral physiological signals

During Fixation, Atomoxetine and Donepezil both reduced posterior cortical alpha-band power relative to Placebo in both the 8-12 Hz (Fig 6A; *p* < 0.05 for all clusters; two-sided cluster-based permutation test) as well as the 2-8 Hz frequency ranges (Fig S7A). This suppression in low-frequency power under cholinergic boost is consistent with previous work in rodents [17,18] and humans [43].

**Fig 6.**
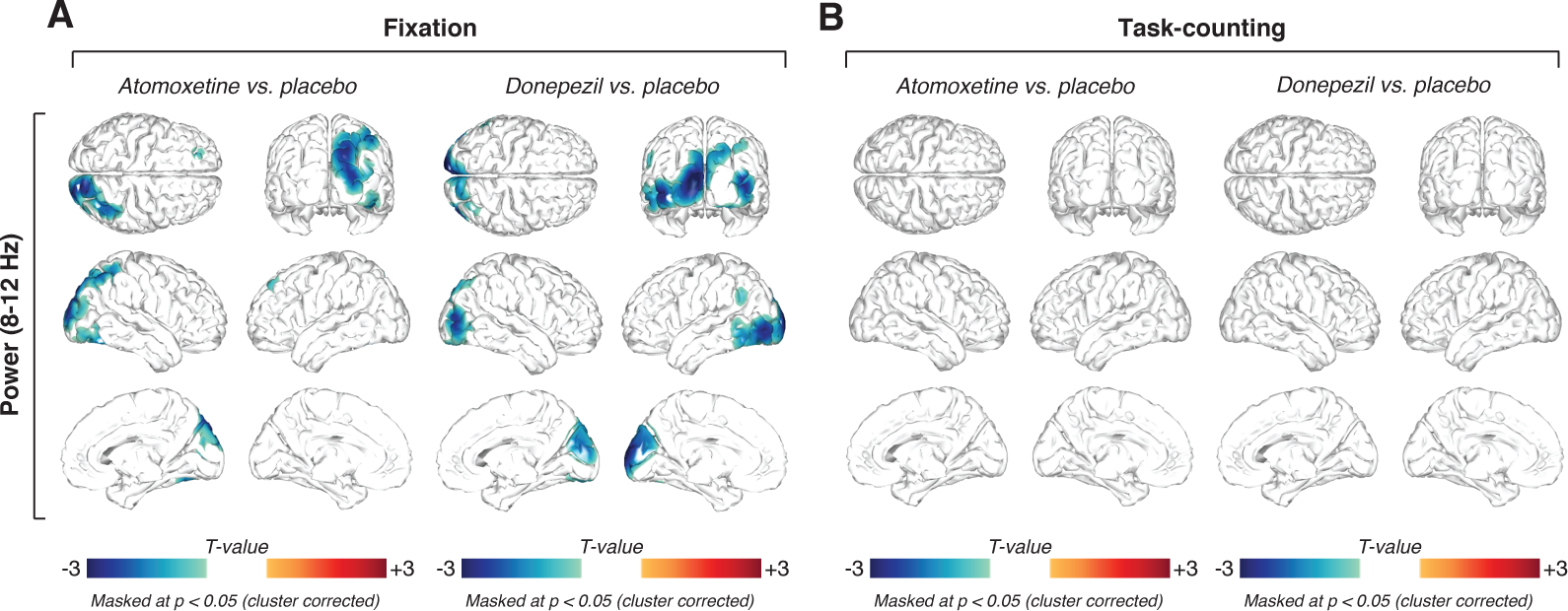
Similar effects of Atomoxetine and Donepezil on 8-12 Hz power. **(A)** Spatial distribution of drug-related alpha power changes during Fixation, thresholded at *p* = 0.05 (two-sided cluster-based permutation test). *Left*. Power changes after the administration of atomoxetine. *Right*. Power changes after the administration of donepezil. **(B)** Same as (A), but for Task-counting. All threshold at *p* = 0.05, cluster-based two-sided permutation tests (N=28).

The Atomoxetine-induced changes on 8-12 Hz power exhibited a different spatial pattern from the one of corresponding change in the scaling exponent *α*: within the cluster of the significant main effect of Atomoxetine on *α* (Fig 4D), power did not correlate with the changes in *α* (group average spatial correlation between pooled difference maps within cluster; *r* = 0.073; *p* = 0.129, *BF* = 1.065). During Task-counting, neither drug altered MEG-power in the low-frequencies (8-12 Hz: Fig 6B, p > 0.05 for all clusters; two-sided cluster-based permutation test; 2-8 Hz: Fig S7B), presumably due to the already suppressed power in the 8-12 Hz range in that condition (Fig 1C). Together with the findings reported in the previous section, the analyses of the mean MEG power indicate that (i) both drugs reduced the amplitude of cortical low-frequency oscillations and (ii) MEG power and the scaling exponent *α* reflected at least partially distinct aspects of intrinsic cortical dynamics

We also controlled for changes in peripheral physiological signals under the drugs as potential confounds of the effect on cortical scaling behavior (Fig 7).As expected, Atomoxetine increased average heart rate (Fig 7A,B). Donepezil had no detectable effect on average heart rate, during neither Fixation (*p* = 0.8676; *t* = 0.16; paired t-test; *BF* = 0.8676; Fig 7A) nor Task-counting (*p* = 0.3274; *t* = 1.0; paired t-test; *BF* = 0.3139; Fig 7B). Both drugs altered heart-rate variability, increasing *α* computed on the time series of inter-heartbeat-intervals (see Methods) in both behavioral contexts relative to Placebo (Fixation: *p* = 0.0012, *t* = 3.62; Task-counting: *p* = 0.0167; *t* = 2.55; Fig 7C; Fixation/Donepezil: *p* = 0.0076, *t* = 2.88; Task-counting/Donepezil: *p* = 0.0049, *t* = 3.06; Fig 7D; all paired t-tests). Critically, the Atomoxetine-induced changes in heart rate showed no (Task-counting: *r* = 0.00; *p* = 0.99; Person correlation; *BF* = 0.15) or only weak and statistically non-significant (Fixation: *r* = 0.24; *p* = 0.21; Person correlation; *BF* = 0.31) correlations with the changes in cortical activity (Fig 7A/B, right). Similarly, the Atomoxetine-related changes in the scaling behavior of inter-heartbeat intervals were not correlated with the changes in cortical scaling behavior (Fixation: *r* = 0.22; *p* = 0.26; *BF* = 0.27; Task-counting: *r* = 0.26; *p* = 0.19; *BF* = 0.35; Fig 7C/D, right). Atomoxetine also decreased spontaneous blink rate during Fixation (*p* = 0.034; *t* = 2.24; paired t-test), but not during Task-counting (*p* = 0.112; *t* = 1.645; *BF* = 1.130; paired t-test; Fig S2B). However, again there was no significant correlation between changes in blink-rate and changes in cortical scaling behavior due to Atomoxetine (Fixation: *r* = −0.26; *p* = 0. 19; *BF* = 0.35; Task-counting: *r* = −0.09; *p* = 0.64; *BF* = 0.16).

**Fig 7.**
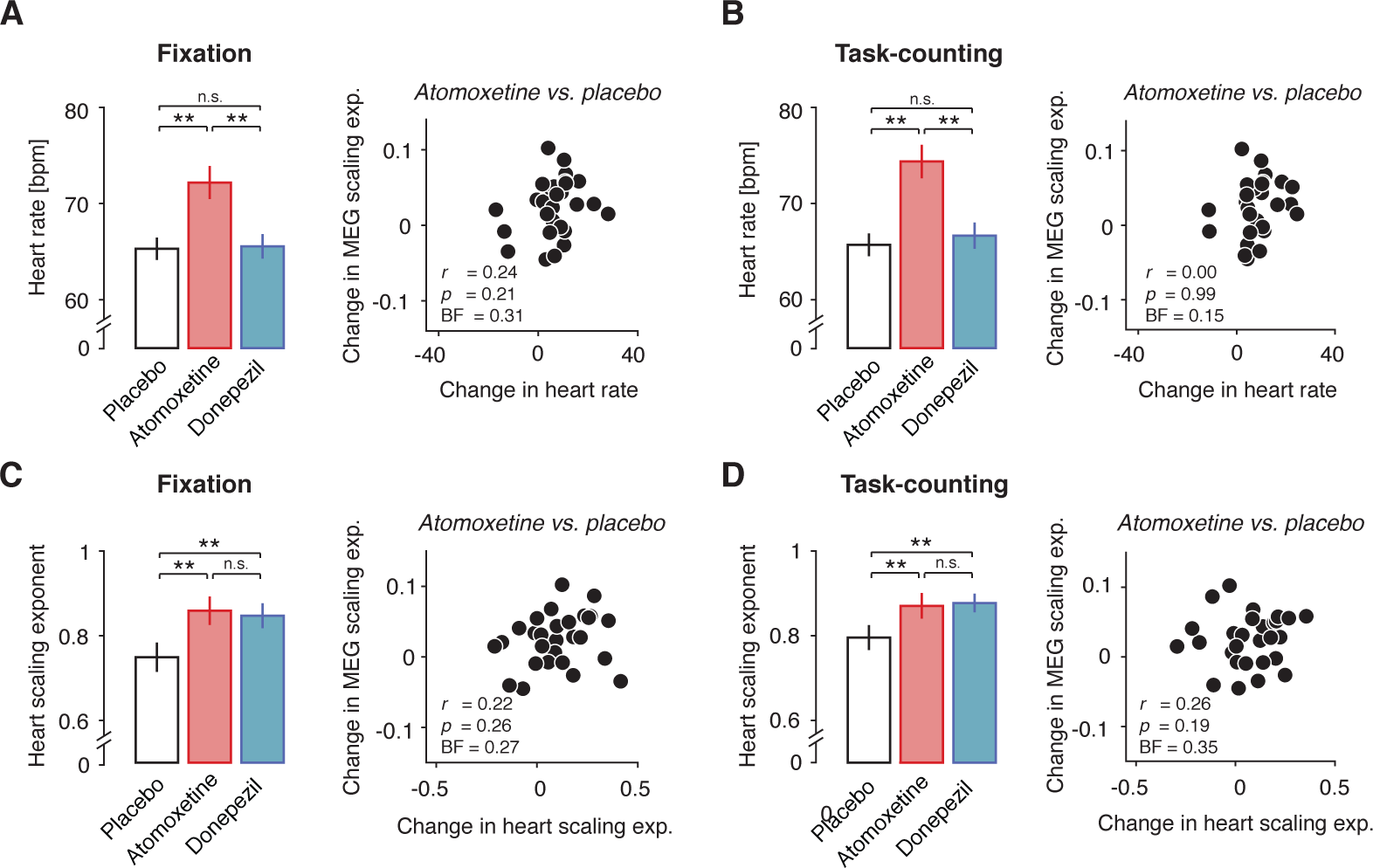
Drug effect on cortical scaling behavior is not explained by systemic drug effects. **(A)** *Left*. Heart rate for Atomoxetine, Placebo and Donepezil during Fixation. *Right*. Correlation of Atomoxetine-related changes in heart rate (x-axis) with Atomoxetine-related changes in MEG scaling exponent *α* (y-axis) (within significant cluster during Fixation). **(B)** As (A), but during Task-counting **(C)** *Right*. Scaling behavior of inter-heartbeat intervals (heart scaling exponent). *Left*. Heart scaling exponent for all pharmacological conditions during Fixation. *Right*. Correlation of Atomoxetine-related changes in heart scaling exponent (x-axis) with Atomoxetine-related changes in MEG scaling exponent *α* (y-axis). **(D)** Same as (C), but during Task-counting. Two-sided t-tests and Pearson correlations (N=28). BF, Bayes factor.

In sum, drug-induced changes in peripheral physiological signals under the drugs, if present, did not account for the Atomoxetine-induced changes in the scaling behavior of the fluctuations in cortical activity (Figs 4 and 5). These controls support our interpretation in terms of a specific effect on cortical net E/I ratio rather than non-specific secondary effects due to the systemic drug effects or changes in retinal input due to blinks.

### Change in scaling exponent under Atomoxetine is consistent with increase in net E/I ratio in cortical circuits

Atomoxetine had an effect on perceptual fluctuations that was in line with a relative increase in excitation in cortical circuits of occipital and posterior parietal cortex that processed the ambiguous visual motion stimulus. We reasoned that this change in circuit state might have also produced the observed change in scaling behavior of intrinsic cortical activity fluctuations under Atomoxetine. In order to solidify this intuition, we simulated the activity of a neural network model made up of recurrently connected excitatory and inhibitory integrate-and-fire units (Fig 8). In what follows, we use the term <u>“E/I ratio”</u> to refer to the ratio of excitatory and inhibitory activity across the circuit [44] and “E/I balance” to refer to a specific regime of E/I ratios, in which excitation and inhibition changes in a proportional way [45–48].

**Fig 8.**
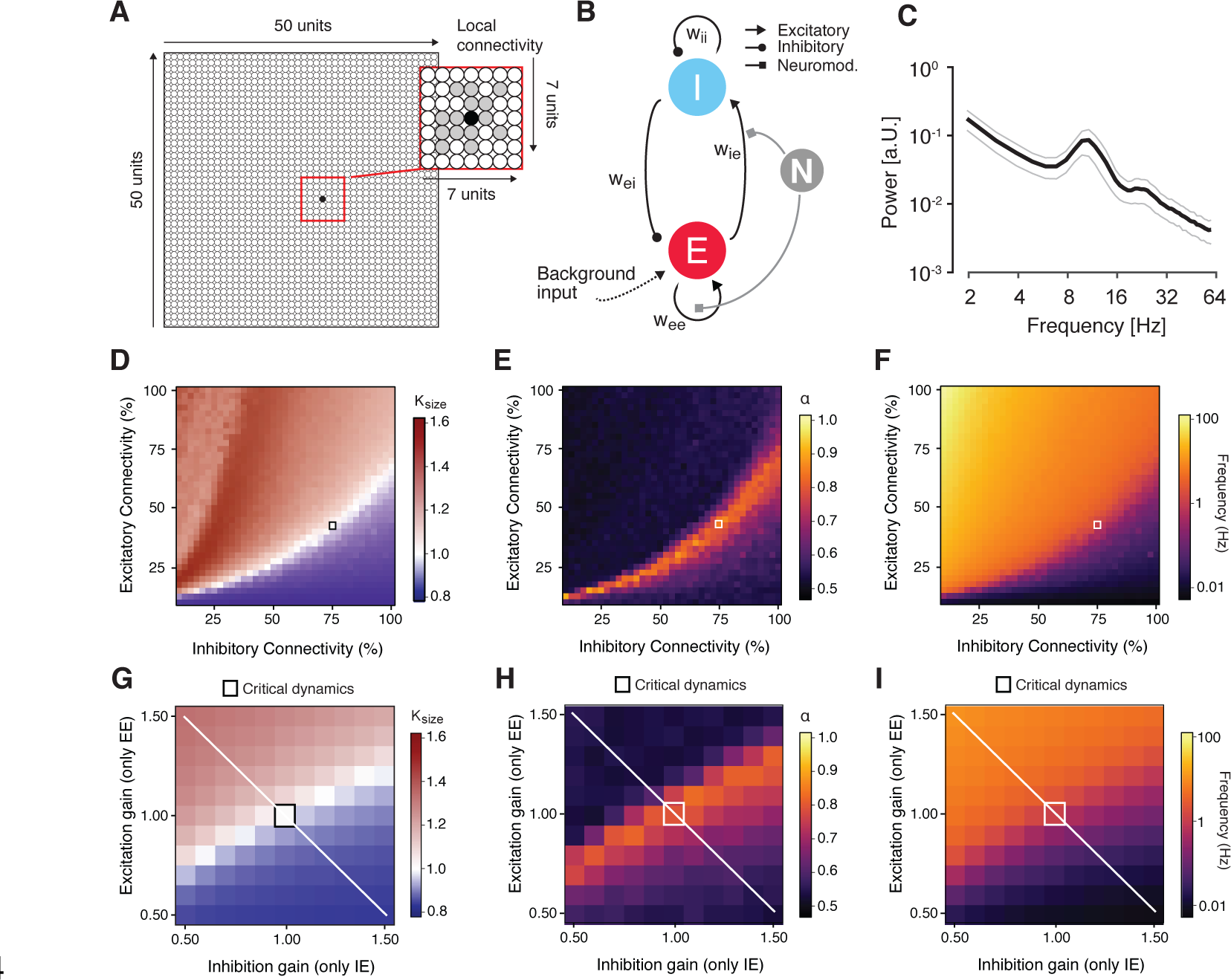
Changes in scaling behavior in a neural network under gain modulation. **(A)-(I)** Dynamic modulation of excitation-inhibition ratio alters long-range temporal correlations in recurrent network model. **(A)** Model architecture. The network consisted of 2500 excitatory and inhibitory integrate-and-fire units and random, local (within an area of 7×7 units) connectivity (magnified within the red square). **(B)** Neuromodulation was simulated as a gain modulation term multiplied with excitatory synaptic weights (*w_ee_* and *w_ie_*). **(C)** Power spectrum of the simulated neural mass activity, with a peak in the alpha range. **(D)** *κ* as a function of excitatory and inhibitory connectivity (with a spacing of 2.5%; means across 10 simulations per cell). The region of *κ*~1, overlaps with the region of *α* > 0.5 and splits the phase space into an excitation-dominant (*κ*>1) and an inhibition-dominant region (*κ*<1). The black square depicts the network configuration that was chosen for assessing the effects of neuromodulation **(E)** Scaling exponent *α* as a function of excitatory and inhibitory connectivity. **(F)** Same as (D) and (E), but for mean firing rate. **(G)** *κ* as a function of independent synaptic gain modulation. Red square, baseline state of critical network before synaptic gain modulation. White line, axis corresponding to largest change in ratio **(H)** Same as (D), but for scaling exponent α. **(I)** Same as (G) and (H), but for firing rate.

We started from a network (Figure 8A) that was similar to the one developed and analyzed in a previous study [49]. The basic features of the model were as follows. The model was built to generate oscillations of neural mass activity (summed across all units) in the alpha-band (8-12 Hz; Figure 8B). The amplitude envelope of these oscillations fluctuated over time, with scale-free long-range temporal correlations. Those scale-free intrinsic fluctuations in cortical activity were sensitive to variations in the percentage of excitatory and inhibitory connections in the circuit (i.e., microstructure). Our previous work [49], reproduced here (Figure 8D-F) showed that such a model accounts for the joint emergence of two scale-free phenomena at different spatial scales (single unit activity vs. mass activity) and temporal scales (tens of milliseconds vs. hundreds of seconds): (i) neuronal avalanches with an event size distribution following a power-law; and (ii) long-range temporal correlations of the amplitude envelope fluctuations of the circuits mass activity. Both phenomena have been established in empirical measurements of cortical population activity [26,50]. Neuronal avalanches are activity deflections (i.e., exceeding a certain threshold) that propagate through the cortical network [50], with an “event size” corresponding to the number of activated units. In line with (Shew et al., 2009) we quantified the power-law scaling of the size distributions of avalanches in the model with the kappa-index (κ): the similarity between the actual event size distribution and a power-law distribution with an exponent of −1.5; A *κ* of 1 indicates perfect match between the two.

We extended this model by means of a multiplicative modulation of synaptic gain [37,51] (Fig 8B). This allowed us to explore how catecholaminergic effects on neural circuits might change the two phenomena of scale-free neural population activity described above. We first determined the structural connectivity (small squares in Fig 8D-F) and the time scale parameters of the model such that the network generated intrinsic alpha-band oscillations (Fig 8C) with amplitude fluctuations that exhibited neuronal avalanches with scale-free event size distributions (Fig 8D) as well as long-range temporal correlations (with *α* ~ 0.85). We then independently modulated specific excitatory or inhibitory connections through the multiplicative scaling of the corresponding synaptic weights, in two ways. In the version shown in Fig 8, we modulated only excitatory synapses, but independently on excitatory as well as inhibitory neurons (EE and IE), thus producing asymmetries in the circuits net E/I ratio as in recent modeling work on a cortical circuit for perceptual decision-making [44]. In the second version (Fig S8A), we co-modulated EE and IE and independently modulated inhibitory synapses on excitatory neurons (EI). This was intended to specifically simulate glutamate receptors (AMPA or NMDA) in former two cases (mediating the effects of excitatory neurons) as opposed to modulations of GABA receptors in the latter case (mediating the effects of inhibitory neurons on others). *N_EE_* and *N_IE_* were co-modulated by the same factor for simplicity, because we did not assume that excitatory (glutamatergic) synapses would be differentially modulated depending on whether they were situated on excitatory or inhibitory target neurons.

Both types of changes in net E/I ratio robustly altered *κ* (Fig 8G and Fig S8B), *α* (Fig 8H and Fig S8C), and the mean firing rate (Fig 8I). The effect of changes in E/I ratio on the scaling exponent *α* were non-monotonic, dependent on the starting point: increases in excitation led to increases in *α* when starting from an inhibition-dominant point, but to decreases in *α* when starting from an excitation-dominant point (Fig 8H, white line). The effects of excitatory and inhibitory gain modulation on the temporal correlation structure of the simulated population activity were qualitatively similar to the effects of changes in the fraction of excitatory and inhibitory synapses simulated (as shown in Fig 8D-F). The latter simulated individual differences in cortical anatomical microstructure, and the former simulated state-dependent changes in cortical circuit interaction, which occur within an individual brain.

In the model, the scaling exponent *α* exhibited a non-monotonic dependence on E/I ratio (see the white diagonal line in Fig 8G-I and schematic depiction in Fig 9). Consequently, without knowing the baseline state, any change in *α* was ambiguous with respect to the direction of the change in E/I ratio (i.e., towards excitation- or inhibition-dominance). Thus, the observed increase in *α* under Atomoxetine during Fixation could have been due to either an increase or a decrease in E/I ratio.

**Fig 9.**
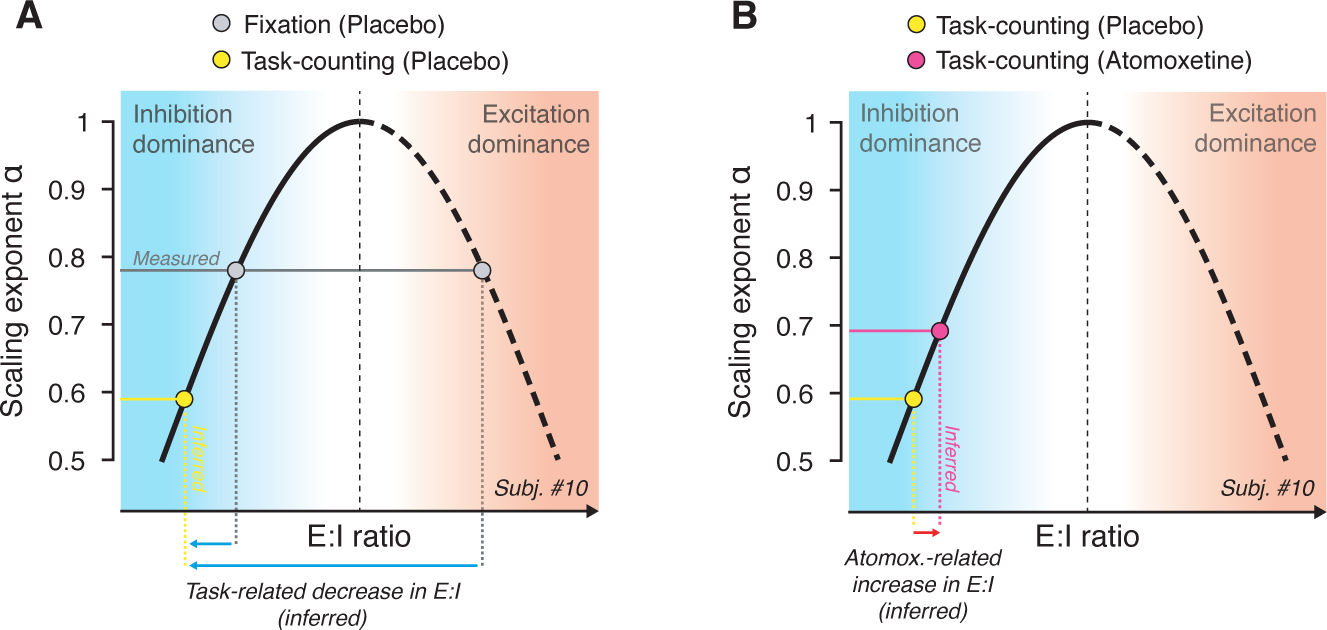
Schematic of inference from observed change in scaling exponents to net E/I ratio (see Results for details). The non-monotonic dependence of scaling exponent *α* on E/I ratio (corresponding to white line in panel H) is replotted schematically. **(A)** Measured scaling exponent *α* during Fixation (gray) can result from both, inhibition- or excitation-dominant regimes; the baseline is unknown. Assuming that sensory drive (Task-counting; yellow dot) either decreases or does not change E/I ratio, the observed decrease in scaling exponent during Task-counting (yellow) reflects a shift towards the inhibition-dominance (blue arrows), consistent with animal physiology [52,53]. **(B)** This constrains the baseline state for the interpretation of the Atomoxetine-induced increase in scaling exponent during Task-counting (red): The latter increase likely reflects an increase in E/I ratio (red arrow).

Importantly, insights from animal physiology helped constrain the baseline state during Task-counting: In the awake state, visual drive decreases E/I ratio in visual cortex V1, due to the recruitment of inhibitory mechanisms that outweigh the excitatory sensory drive [52,53]. We assumed that the same held for the Task-counting condition (constant visual stimulation) of our study.

This condition enabled us to infer the change in net E/I ratio under Atomoxetine. The rationale is illustrated in Fig 9. The animal physiology results referred to above indicte that the observed decrease in *α* during Task-counting was due to a shift towards inhibition-dominance (yellow point in Fig 9A). Under this assumption, the Atomoxetine-induced increase in *α* during was due to an increase in net E/I ratio (Fig 9B). Because the effects of Atomoxetine on *α* were the same during Task-counting and Fixation, it is likely that the same mechanism was at play during Fixation.

In sum, under certain, the simulations provided a mechanistic explanation for the observed MEG effects: effective changes in the cortical E/I ratio, due to multiplicative changes of synaptic gain [37] or other mechanisms [12,20] – the same conclusion inferred from the increase in the rate of perceptual alternations above.

## DISCUSSION

Neuromodulators regulate ongoing changes in the operating mode of cognitive processes [1,5,6,10,54] as well as of cortical microcircuits [11,12,17,18,20,21]. Here, we unraveled the effect of two major classes of neuromodulators, catecholamines and acetylcholine, on the intrinsic variability of cortical computation, an important parameter shaping the operating mode. We used two separate read-outs of this parameter: (i) the rate of fluctuations in perception induced by ambiguous visual input and (ii) the temporal correlation structure of (scale-free) fluctuations of the amplitude envelope of band-limited cortical mass activity. Catecholamines, but not acetylcholine, altered both read-outs.

Simulations of a recurrent neural network revealed that, under well supported physiological assumptions, the observed changes in the temporal structure of fluctuations in cortical activity is indicative of an increase in the cortical E/I ratio. Earlier modeling and empirical data show that such an increase in net E/I ratio in visual cortex is also consistent with the increased rate of perceptual fluctuations under the catecholaminergic boost.

### Cortical distribution of Atomoxetine effects on cortical activity fluctuations

The Atomoxetine effects on the scaling exponent were widespread across cortex, but not entirely homogenous. They were pronounced across occipital and parietal cortex, but not robust in frontal cortex (see Fig 5B). This distribution might point to a noradrenergic, rather than dopaminergic origin. Atomoxetine increases the levels of both catecholamines, noradrenaline and dopamine, but the cortical projection zones differ substantially between both systems: Dopaminergic projections mainly target prefrontal cortex [55] and only sparsely to occipital cortex [56,57], whereas the noradrenergic projections are more widespread and strong to occipital and parietal cortex [58]. Alternatively, this distribution may reflect the different receptor composition across cortical regions [58,59]. The relative frequency of the different noradrenaline receptors differs between prefrontal and posterior cortex [58], which might translate a homogenous noradrenaline release into a heterogeneous effect on the activity in these different cortical regions. An important next step will be to investigate the differential role of different noradrenaline receptors, and different regional receptor profiles, in shaping the cortex-wide effect of noradrenaline on long-range temporal correlations.

### Opposite effects of external drive and catecholamines on long-range temporal correlations

Consistent with our current results, previous studies also found a decrease in temporal autocorrelations of cortical activity due to external drive [27,42]. The observation is consistent with the insight from intracellular recordings of cortical neurons in animals, that cortical responses to sensory stimulation in the awake state are dominated by inhibition [52,53,60,61]. One candidate source of this sensory-driven state change is thalamocortical inhibition [62], but intracortical feedback inhibition might also contribute [63]. Modeling work shows that the driven state is associated with shortened temporal autocorrelations as well as a decrease in the entropy of activity states in large-scale cortical networks [64]. Correspondingly, the increase in long-range temporal autocorrelations under catecholaminergic modulation may be associated with an increase in entropy – that is, a tendency to explore a larger set of cortical activity states. It is tempting to link this to the prominent idea that high sustained noradrenaline levels promote an exploratory mode of cortical computation and behavior [5].

### Convergent evidence for catecholaminergic increase in cortical E/I ratio

Cortical circuits maintain a tight balance between excitation and inhibition, which is largely preserved across contexts and levels of the cortical hierarchy [46,48]. However, even in the absence of changes in sensory input, neuromodulators such as noradrenaline and acetylcholine can change the cortical E/I ratio [65,66]. The E/I ratio, in turn, shapes the computational properties of cortical circuits [67,68], and thereby the behavior of the organism [37,44,69]. Substantial evidence already points to significant changes in E/I ratio in schizophrenia and autism [70–72]. Similar changes might be at play in other brain disorders as well [73].

Our simulations indicated that the temporal correlation structure of neural population activity, as measured with the scaling exponent *α*, is sensitive to changes in E/I ratio, produced through synaptic gain modulation (see the white line in Fig 8H). In both versions of our model, the neuromodulatory effects were not perfectly symmetric (see the deviations of peak scaling exponents from main diagonal in Fig 8H). While the latter effect was small and may be specific to the details of the model, it remains possible that the subtle changes in scaling exponents we observed were produced through symmetric gain modulations that maintained the net E/I balance (i.e., along the main diagonal). However, two additional lines of evidence converge on our conclusion that catecholamines (in particular noradrenaline) boosted E/I ratio. First, in the same participants, the catecholaminergic manipulation had a reliable effect on the perceptual switch rate, which is also indicative of cortical E/I ratio [33,34,36]. Second, results invasive rodent work also point to an increase in cortical E/I ratio under noradrenaline: Noradrenaline was found to decrease spontaneous inhibition in auditory cortex [66] and mediate a tonic depolarization of visual cortical neurons during locomotion [20].

### No evidence for donepezil effects on cortical or perceptual fluctuations

The absence of an effect of Donepezil on either perceptual fluctuations or long-range temporal correlations of cortical activity may be due to the small dosage or the single administration of the drug in our study. Even so, our donepezil manipulation was sufficient to robustly change heart rate variability and, more importantly, low-frequency power of cortical activity, an established marker of cholinergic action in cortex [17,18,43,74]. The lack of the effects of donepezil on perceptual fluctuations and cortical scaling behavior might also be due to the specific properties of cholinergic action on the cortical net E/I ratio. Invasive evidence indicates that acetylcholine can rapidly disinhibit pyramidal cells by activating a chain of two inhibitory interneurons [21], a mechanism that may alter E/I ratio mainly during stimulus-evoked responses [65]. By contrast, noradrenaline also alters the levels of tonic inhibition of pyramidal cells occurring spontaneously [66]. This might explain the dissociation between the effects of atomoxetine and donepezil under the current steady-state conditions, which excluded (or minimized) stimulus-evoked transients.

### Functional consequences of changes in net cortical E/I ratio

We observed a selective increase in the rate of spontaneous perceptual alternations under catecholaminergic adding to evidence that these dynamics are under neuromodulatory control [75]. Such a change could be due to an increase in cortical “noise” [33]. Future invasive studies should relate chatecholaminergic changes in the variability of the spiking activity [76] of neurons contributing directly to the contents of multi-stablke perception.

We suspect that an increase in cortical E/I ratio will have particularly strong effects on behavior when affecting parietal and prefrontal cortical circuits characterized by slow intrinsic timescales [30,32,77] and involved in persistent activity during working memory and the slow accumulation of information over time [69]. It is possible that the catecholaminergic effects on parietal cortex we observed here reflects an increase in the recurrent excitation, which, is essential for sustained processes such as working memory [78] as well as information integration during decision-making [32,79]. Future work should assess this through the use of tasks probing into network reverberation and information accumulation in association cortex.

### A control parameter for critical network dynamics

In our model, long-range temporal correlations in the fluctuations of neural mass activity (i.e., activity summed across the entire local network) [26] and avalanches within the neuronal network [50] jointly emerge at the same E/I ratio. Both phenomena are commonly interpreted as hallmarks of “criticality” [26,29,50,80] – a state of a complex dynamical system poised between order and chaos [81–83]. It has been proposed that the cortex operates in a narrow regime around criticality [83,84], potentially optimizing its computational capacities [80,85–88]. A number of reports showed that cortical dynamics may continuously vary around the critical state [89–92], but the source of these fluctuations has, so far, remained unknown. Here, we have identified catecholaminergic neuromodulation as an endogenous factor controlling these spontaneous variations in critical dynamics.

In complex systems, critical dynamics can emerge in a self-organized fashion [81], or through an external control parameter that fine-tunes the system. The tuning of temperature in the Ising model of spin magnetization [83] is a common example for the latter case. It is tempting to speculate that catecholaminergic tone serves as such a control parameter in the cerebral cortex.

### Link between catecholaminergic effects on fluctuations in perception and cortical mass activity

We here used two read-outs of catecholaminergic effects, constituting two distinct expressions of the resulting changes in cortical circuit state. The envelope of cortical alpha-band oscillations collapsed across large chunks of cortex is unlikely to encode the contents of perception in the phenomenon studied here. The perceived direction of 3D-motion, which fluctuates spontaneously, is encoded in fine-grained patterns of neural population activity within motion-sensitive visual cortical areas [39,93]. The power of alpha-band oscillations is a more global feature of cortical population activity, which is likely insensitive to the fine-grained, within-area patterns of neural population activity. The widespread release of neuromodulators changes also the cortical circuit state, specifically E/I ratio, in a widespread manner. Such changes, in turn, alter the highly specific (fine-grained) interactions between percept-selective populations of visual cortical neurons that give rise to the perceptual dynamics [33,34,36]. Thus, although both read-outs likely tap into similar changes in global cortical circuit state, there is no one-to-to mapping between them.

### Limitations of the current modeling approach

While our model simulations provided important mechanistic insights, the model has limitations that should be addressed in future work. First, different from the MEG data, the power of alpha-band oscillations behaves similarly to the scaling exponents in the model (Fig S8E). This is because the model oscillations emerge from the same recurrent neuronal interactions within the patch that also shape the long-range temporal correlations in the envelopes of the amplitude envelopes of these oscillations. By contrast, in the brain, alpha-band power of local cortical mass signals is likely affected by a variety of sources other than local circuits, for instance alpha-frequency modulated input from the thalamus [94]. This might lead to dissociations between changes in MEG power and long-range temporal correlations of the power fluctuations which the model does not capture in its present form. Second, the model lacks long-range excitatory connections, which are prominent in the real cortex, and whose effects on the correlation structure of cortical fluctuations are largely unknown.

### Conclusion

The combined measurement of fluctuations in bistable perception as well as in cortical mass activity under steady-state conditions provides an easily assessable, multi-level read-out of pharmacological effects on cortical computation. In our study, this read-out supported the idea that catecholamines boost the intrinsic variability of perception and behavior, an effect that might be mediated by an increase in the net E/I ratio in the visual cortical system. This read-out may be useful for inferring changes in cortical E/I ratio in neuropsychiatric disorders, or in their pharmacological treatment in future work.

## METHODS

### Pharmacological MEG experiment

#### Participants

30 healthy human participants (16 females, age range 20-36, mean 26.7) participated in the study after informed consent. The study was approved by the Ethical Committee responsible for the University Medical Center Hamburg-Eppendorf. Two participants were excluded from analyses, one due to excessive MEG artifacts, the other due to not completing all 3 recording sessions. Thus, we report results from N=28 participants (15 females). In one of those participants, one Task-counting run (during the Atomoxetine condition) was not recorded due to a software problem with the data acquisition computer.

#### General design

We pharmacologically manipulated the levels of catecholamines (noradrenaline and dopamine) and acetylcholine in a double-blind, randomized, placebocontrolled, and cross-over experimental design (Fig 1A, B). Each participant completed three experimental sessions, consisting of drug (or placebo) intake at two time points, a waiting period of 3 hours, and an MEG recording. During each MEG session, participants were seated on a chair inside a magnetically shielded MEG chamber. Each session consisted of 6 runs of different tasks, each of which was 10 minutes long and followed by breaks of variable duration.

#### Pharmacological intervention

We used the selective noradrenaline reuptake inhibitor atomoxetine (dose: 40 mg) to boost the levels of catecholamines, specifically noradrenaline and (in prefrontal cortex) dopamine [10]. We used the cholinesterase inhibitor donepezil (dose: 5 mg) to boost acetylcholine levels. Atomoxetine is a relatively selective inhibitor of the noradrenaline transporter, which is responsible for the natural reuptake of noradrenaline that has been released into the extracellular space. Consequently, atomoxetine acts to increase the extracellular levels of noradrenaline, an effect that has been confirmed experimentally in rats prefrontal cortex [95]. The same study showed that atomoxetine also increases the prefrontal levels of dopamine, which has a molecular structure very similar to the one of noradrenaline and is, in fact, a direct precursor of noradrenaline. Atomoxetine has smaller affinity to the serotonin transporter, and there are discrepant reports about the quantitative relevance of these effects: While one study found no increases in serotonin levels under atomoxetine [95], a recent study reports a significant atomoxetine-related occupancy of the serotonin transporter in non-human primates [96] at dosages which would correspond to human dosages of 1.0 – 1.8 mg/kg. Note, that these dosages are substantially higher than the administered dosage in this study (40 mg, independent of body weight). It is, therefore, unclear to which extent our Atomoxetine condition affected cortical serotonin levels.

Donepezil is a selective inhibitor of the enzyme acetylcholinesterase, which breaks up all the extracellular acetylcholine to terminate its synaptic action. Consequently, donepezil acts to increase the extracellular levels of acetylcholine. Donepezil is also an agonist of the endoplasmatic sigma1-receptor, which modulates intracellular calcium signaling.

A mannitol-aerosil mixture was administered as placebo. All substances were encapsulated identically in order to render them visually indistinguishable. Peak plasma concentrations are reached ~3-4 hours after administration for donepezil [97] and 1-2 hours after administration for atomoxetine [98], respectively. We adopted the following procedure to account for these different pharmacokinetics (Fig 1A): participants received two pills in each session, one 3 h and another 1.5 h before the start of MEG recording. In the Atomoxetine condition, they first received a placebo pill (t = −3 h) followed by the atomoxetine pill (t = −1.5 h). In the Donepezil condition, they first received the donepezil pill (t = −3 h), followed by placebo (t = −1.5 h). In the Placebo condition, they received a placebo at both time points. The half-life is ~ 5 h for atomoxetine [98] and ~ 82 h for donepezil, respectively [97]. In order to allow plasma concentration levels to return to baseline, the three recording sessions were scheduled at least 2 weeks apart. This design ensured maximum efficacy of both pharmacological manipulations, while effectively blinding participants as well as experimenters.

#### Stimuli and behavioral tasks

In each session, participants alternated between three different task conditions (2 runs à 10 minutes per condition) referred to as Fixation, Task-counting, and Task-pressing in the following (Fig 1B). All conditions entailed overall constant sensory input. Fixation and Task-counting also entailed no overt motor responses and are, therefore, referred to as “steady-state” conditions in the following. We used these steady-state conditions to quantify intrinsic fluctuations in cortical activity. Task-pressing entailed motor responses and was used for reliable quantification of perceptual dynamics. All instructions and stimuli were projected onto a screen (distance: 60 cm) inside the MEG chamber. The individual conditions are described as follows.

##### Fixation

Participants were asked to keep their eyes open and fixate a green fixation dot (radius = 0.45° visual angle) presented in the center of an otherwise gray screen. This is analogous to eyes-open measurements of “resting-state” activity widely used in the literature on intrinsic cortical activity fluctuations.

##### Task-counting

Participants viewed a seemingly rotating sphere giving rise to the kinetic depth effect [99,100]: spontaneous changes in the perceived rotation direction (Fig 1B). The stimulus subtended 21° of visual angle. It consisted of 1000 dots (500 black and 500 white dots, radius: 0.18° of visual angle) arranged on a circular aperture presented on a mean-luminance gray background, with the green fixation dot in the center. In order to minimize tracking eye movements, the sphere rotation was along the horizontal axis, either “forward” (towards the observer) or “backward” (away from the observer), and the dot density decreased along the horizontal axis towards the center of the stimulus. Participants were instructed to count the number of perceived changes in rotation direction and report the total number of perceived transitions at the end of the run. Just like during Fixation, Task-counting minimized any external (sensory or motor) transients. Subjects silently counted the alternations in perceived rotation direction and verbally reported the total count after the end of the 10 minutes run.

##### Task-pressing

This condition was identical to Task-counting, except that participants were instructed to press and hold one of two buttons with their index finger to indicate the perceived rotation direction of the sphere. Thus, each perceptual alternation was accompanied by a motor response leading to change in the button state. This allowed for a more reliable quantification of participants’ perceptual dynamics. On two sessions (Atomoxetine condition), button presses were not registered. Hence, the corresponding analyses were performed on 26 participants.

### Data acquisition

MEG was recorded using a whole-head CTF 275 MEG system (CTF Systems, Inc., Canada) at a sampling rate of 1200 Hz. In addition, eye movements and pupil diameter were recorded with an MEG-compatible EyeLink 1000 Long Range Mount system (SR Research, Osgoode, ON, Canada) at a sampling rate of 1000 Hz. In addition, electrocardiogram (ECG) as well as vertical, horizontal and radial EOG were acquired using Ag/AgCl electrodes (sampling rate 1200 Hz).

### Data analysis

#### Eye data

Eye blinks were detected using the manufacturer’s standard algorithm with default settings. Saccades and microsaccades were detected using the saccade detection algorithm described in [101], with a minimum saccade duration of 4 samples (= 4 ms) and a threshold velocity of 6. For 18 out of 28 participants, only horizontal eye movements were recorded.

#### EOG data

EOG events (blinks and saccades) were extracted using semi-automatic artifact procedures as implemented in FieldTrip [102]. In short, EOG traces were bandpass filtered using a third-order butterworth filter (1 – 15 Hz) and the resulting signal was z-scored. All time points where the resulting signal exceeded a z-score of 4 were marked as an EOG event.

#### MEG data

##### Preprocessing

First, all data were cleaned of strong transient muscle artifacts and squid jumps through visual inspection and manual as well as semi-automatic artifact rejection procedures, as implemented in the FieldTrip toolbox for MATLAB [102]. To this end, data segments contaminated by such artifacts (+/− 500 ms) were discarded from the data (across all channels). Subsequently, data were downsampled to 400 Hz split into low (2-40 Hz) and high (>40 Hz) frequency components, using a 4th order (low- or high-pass) Butterworth filter. Both signal components were separately submitted to independent component analysis [103] using the FastICA algorithm [104]. Artifactual components (eye blinks/movements, muscle artifacts, heartbeat and other extra-cranial artifacts) were identified based on three established criteria [105]: power spectrum, fluctuation in signal variance over time (in bins of 1s length), and topography. Artifact components were reconstructed and subtracted from the raw signal and low- and high frequencies were combined into a single data set. On average, 20 (+/− 14) artifact components were identified for the low frequencies and 13 (+/− 7) artifactual components were identified for the high frequencies.

##### Spectral analysis

Sensor-level spectral estimates (power spectra and cross spectral density matrices) were computed by means of the multi taper method using a sequence of discrete prolate Slepian tapers [106]. For the power spectrum shown in Fig 1C, power spectra were computed using a window length of 5s and a frequency smoothing of 2 Hz, yielding 19 orthogonal tapers. The focus of this paper was on the fluctuations of the amplitude envelopes, rather than on the (oscillatory) fluctuations of the carrier signals *per se*. The temporal correlation structure of the amplitude envelope fluctuations of cortical activity seems similar across different carrier frequency bands [29]. We focused on amplitude envelope fluctuations in the alpha-band because (i) the cortical power spectra exhibited a clearly discernible alpha-peak, which robustly modulated with task, as expected from previous work [40] (Fig 1C); and (ii) the computational model used to study the effect of synaptic gain modulation on cortical activity fluctuations was tuned to produce alpha-band oscillations (see above and [49]).

##### Source reconstruction: general approach

The cleaned sensor level signals (*N* sensors) were projected onto a grid consisting of *M* = 3000 voxels covering gray matter of the entire brain (mean distance: 6.3 mm) using the exact low-resolution brain electromagnetic tomography (eLORETA; [107] method. The grid was constructed from the ICBM152 template [108], covering gray matter across the brain. The magnetic leadfield was computed, separately for each subject and session, using a single shell head model constructed from the individual structural MRI scans and the head position relative to the MEG sensors at the beginning of the run [109]. In case no MRI was available (4 subjects), the leadfield was computed from a standard MNI template brain transformed to an estimate of the individual volume conductor using the measured fiducials (located at the nasion, the left and the right ear).

In order to depict the source-level results, we interpolated the voxel-level results onto the surface of the brain. Activations from structures distant to the surface are not shown and were exponentially attenuated.

##### Source level estimates of amplitude envelopes and power

For comparing amplitude envelope and power estimates between experimental conditions in source space we aimed to select a single direction of the spatial filter for each voxel across pharmacological conditions (i.e., MEG sessions), but separately for Fixation and Task-Counting conditions. The rationale was to avoid filter-induced biases in the comparisons between the pharmacological conditions, while allowing that external task drive might systematically change the dipole orientations.

To this end, we first computed the mean source-level cross-spectral density matrix *C*(*r*,*f*) for each frequency band, *f*, averaged across the three MEG sessions, as follows:

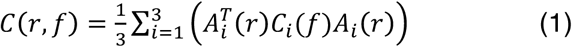

whereby *i* indicated the MEG session, *C_i_*(*f*) was the (sensor-level) session- and frequency-specific cross-spectral density matrix and *A_i_* is the spatial filter for session *i*. We then extracted the first eigenvector *u*_1_(*r*, *f*) of the session-average matrix *C*(*r*,*f*), by means of singular value decomposition, and computed the unbiased filter selective for the dominant dipole orientation, *B_i_*(*r*, *f*), as:

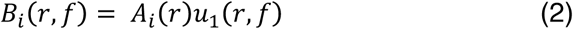

This procedure ensures that, for each voxel, dipole orientation was chosen such that power is maximized. Please note that this filter was now frequency-specific, whereas the previous filters, *A_i_*(*r*)), were not. To obtain instantaneous estimates of source-level amplitudes, the sensor-level signal for session *i*, *X_i_*(*t*), was band-pass filtered (using a finite impulse response filter) and Hilbert-transformed, yielding a complex-valued signal *H_i_*(*f*,*t*) for each frequency band. This signal was projected into source space through multiplication with the unbiased spatial filter, *B_i_*(*r*,*f*), and the absolute value was taken:

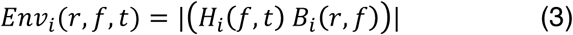

where *Env_i_*(*r*,*f*,*t*) was the estimated amplitude envelope time course of source location *r* and frequency *f*. Next, for each session, unbiased source-level cross spectral density estimates were obtained from the sensor-level cross-spectral density matrix *c_i_*(*f*) and the frequency-specific, unbiased spatial filter *B_i_*(*f*). The main diagonal of the resulting matrix contains source-level power estimates for all source locations:

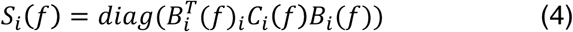

These computations where repeated separately for the Task-counting and Fixation conditions, session by session. The differences in amplitude envelope fluctuations and power estimates between pharmacological and task conditions reported in this paper were robust with respect to the specifics of the analysis approach. In particular, we obtained qualitatively similar pharmacological effects in sensor space, as reported in an earlier conference abstract [110].

##### Detrended fluctuation analysis

The source-level amplitude envelopes *Env_i_*(*r*,*f*,*t*) were submitted to detrended fluctuation analysis [111,112] in order to quantify long-range temporal correlations. Detrended fluctuation analysis quantifies the power law scaling of the fluctuation (root-mean-square) of a locally detrended, cumulative signal with time-window length. Different from the analysis of the more widely known autocorrelation function [30,77], detrended fluctuation analysis provides robust estimates of the autocorrelation structure for stationary and non-stationary time series. The procedure of the detrended fluctuation analysis is illustrated in Fig 2.

For simplicity, in the following, we re-write the amplitude envelope *Env_i_*(*r*,*f*,*t*) as *x* of length *T*. First, we computed the cumulative sum of the demeaned *x*, (Fig 2B):

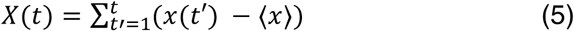

where *t′* and *t* denote single time points up to length *T*. The cumulative signal *X* was then cut into *i* = 1 …*k* segments *Y_i_* of length *N* (overlap: 50%), where *k* = *floor*[(*T* – *N*)/(0.5 *N*)] (Fig 2B, top). Within each segment *Y_i_* of equal length *N*, the linear trend *Y_i_trend_* (least squares fit) was subtracted from *Y_i_* (Fig 2B, bottom, blue vs. red lines), and the root-mean-square fluctuation for a given segment was computed as:

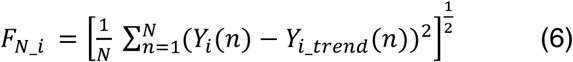

where *n* indicates the individual time points. The fluctuation was computed for all *k* segments of equal length *N* and the average fluctuation was obtained through:

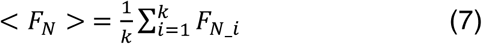

The procedure was repeated for 15 different logarithmically spaced window lengths *N*, ranging from 3 s to 50 s, which yields a fluctuation function (Fig 2C). As expected for scale-free time series (103), this fluctuation function follows a power-law of the form:

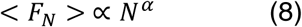

The “scaling exponent” *α* was computed through a linear regression fit in log-log coordinates (Fig 2C). The longest and shortest window lengths were chosen according to guidelines provided in [112].

A scaling exponent of *α* ~= 0.5 indicates a temporally uncorrelated (“white noise”) process. Scaling exponents between 0.5 < *α* < 1 are indicative of scalefree behavior and long-range temporal correlations [112], whereas exponents of *α* < 0.5 indicate long-range anti-correlations (“switching behavior”) and *α* > 1 are indicative of an unbounded process [112]. The scaling exponents for alpha-band MEG amplitude envelopes estimated in this study ranged (across experimental conditions, MEG sensors and participants) from 0.40 and 1.04, with 99.4% of all estimates in the range from 0.5 to 1. This is indicative of scale-free behavior and consistent with previous human MEG work [26–29,42,113].

##### Relationship between measures of cortical variability

Scale-free behavior of neural time series has also been quantified via analysis of the power spectrum [24,25]. There is a straightforward relationship between both approaches, which we explain below, to help appreciate our results in the context of these previous studies. The power spectrum of the amplitude envelope of cortical activity is typically well approximated by the power law *p*(*f*) ∝ *f*^−*β*^, where *β* is referred to as the power-law exponent (Fig 2D). For power-law decaying autocorrelations, the relationship between the power-law exponent *β* and the scaling exponent *α* (estimated through DFA) of a time series is:

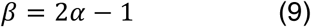

##### Analysis of ECG data

ECG data were used to analyze two measures of peripheral autonomic activity: average heart rate and heart rate variability. For both measures, we used an adaptive threshold to detect the R-peak of each QRS-complex in the ECG. Heart rate was then computed by dividing the total number of R-components by time. Heart rate variability was quantified by means of the detrended fluctuations analysis described for MEG above, but now applied to the time series of the intervals between successive R-peaks [28,29]. In line with the MEG analyses, we used windows ranging from 3 to 50 heartbeats (roughly corresponding to 3–50 s).

#### Statistical tests

Statistical comparisons of all dependent variables between conditions were, unless stated otherwise, performed using paired t-tests.

Null effects are difficult to interpret using regular null hypothesis significance testing. The Bayes Factor addresses this problem by quantifying the strength of the support for the null hypothesis over the alternative hypothesis provided by the data, taking effect size into account. Wherever null effects were conceptually important, results obtained from a regular (paired) t-test [114] and Pearson correlations [115] were converted into corresponding Bayes Factors.

To map significant changes of scaling exponents *α* across the brain, we computed a non-parametric permutation test based on spatial clustering [116,117]. This procedure has been shown to reliably control for Type I errors arising from multiple comparisons. First, a paired t-test was performed to identify voxels with significant changes (voxel with *p* < 0.05). Subsequently, significant voxels are combined into clusters based on their spatial adjacency. Here, a voxel was only included into a cluster when it had at least two significant neighbors. Subsequently, the t-values of all voxels comprising a cluster were summed, which yields a cluster statistic (i.e., a cluster t-value) for each identified cluster.

Next, a randomization null distribution was computed using a permutation procedure (*N* = 10.000 permutations). On each permutation, the experimental labels (i.e., the pharmacological conditions) were randomly re-assigned within participants and the aforementioned procedure was repeated. For each iteration, the maximum cluster statistic was determined and a distribution of maximum cluster statistics was generated. Eventually, the cluster statistic of all empirical clusters was compared to the values obtained from the permutation procedure. All voxels comprising a cluster with a cluster statistic smaller than 2.5% or larger than 97.5% of the permutation distribution were labeled significant, corresponding to a corrected threshold of *α* = 0.05 (two-sided).

### Model simulations

To simulate the effects of synaptic gain modulation on cortical activity fluctuations, we extended a previously described computational model of a local cortical patch [49] by means of multiplicative modulation of synaptic gain. All features of the model were identical to those of the model by [49], unless stated otherwise. The model consisted of 2500 integrate-and-fire neurons (75% excitatory, 25% inhibitory) with local connectivity within a square (width = 7 units) and a connection probability that decayed exponentially with distance (Fig 8A). The dynamics of the units were governed by:

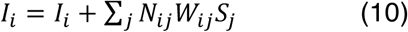

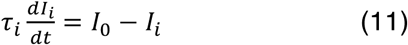

where subscripts *i, j* indicated different units, *N_ij_* was a multiplicative gain factor, *W_ij_* were the connection weights between two units, and *S_j_* a binary spiking vector representing whether unit *j* did or did not spike on the previous time step, and *I*_0_ = 0. For all simulations reported in this paper, we optimized the connection weights using Bonesa [118], a parameter tuning algorithm, such that the network exhibited alpha-band oscillations, long-range temporal correlations, and neuronal avalanches (see below). The optimized values for the connection weights were *W_EE_* = 0.0085, *W_IE_* = 0.0085, *W_EI_* = −0.569 and *w_II_* = −2 whereby subscript *E* indicated excitatory, subscript *I* indicated inhibitory, and the first and second subscript referred to the receiving and sending unit, respectively.

On each time step (*dt* = 1 ms), *I_i_* was updated for each unit *i*, with the summed input from all other (connected) units *j* and scaled by a time constant *τ_i_* = 9 ms, which was the same for excitatory and inhibitory units. The probability of a unit generating a spike output was given by:

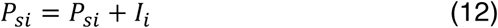

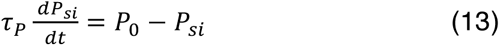

with the time constant for excitatory units *τ_P_* = 6 *ms* and for inhibitory *τ_P_* = 12 *ms*. *P*_0_ was the background spiking probability, with *P*_0_(*exc*.) = 0.000001 [1/*ms*] and *P*_0_(*inh*.) = 0 [1/*ms*]. For each time step, it was determined whether a unit did or did not spike. If it did, the probability of that unit spiking was reset to

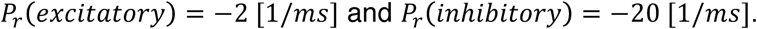

We used this model to analyze the dependency of two quantities on E/I ratio: (i) the power-law scaling of the distributions of the sizes of neuronal avalanches [50] estimated in terms of the kappa-index *κ* which quantifies the difference between an empirically observed event size distribution and a theoretical reference power-law distribution with a power-law exponent −1.5 [86], and (ii) the scaling behavior (scaling exponent *α*) of the amplitude envelope fluctuations of the model’s local field potential. To this end, we summed the activity across all (excitatory and inhibitory) neurons to obtain a proxy of the local field potential. We band-pass filtered the local field potential in the alpha-band (8–12 Hz) and computed long-range temporal correlations in the alpha-band amplitude envelopes following the procedure described above (see *Detrended fluctuation analysis of MEG data*), using windows sizes ranging from 5 s to 30 s.

In order to assess the influence of structural excitatory and inhibitory connectivity on network dynamics (Figs 4D-F), we varied the percentage of units (excitatory and inhibitory) a given excitatory or inhibitory unit connects to within a local area (7 units × 7 units; Fig 8A). These percentages were varied independently for excitatory and inhibitory units with a step size of 2.5%.

The gain factor *N_ij_* was the main difference to the model described by [49]. It was introduced to simulate the effects of neuromodulation on synaptic interactions in the cortical network [37]. For this, we kept all the structural parameters fixed (42.5% excitatory connectivity, 75% inhibitory connectivity; small square in Figs 4D-F), in a range where the model exhibits both robust long-range temporal correlations as well as neuronal avalanches. Note that any other combination of parameters would yield similar results, as long as the model exhibits these two phenomena. From the chosen starting point, we systematically varied the synaptic gain factors, in two different ways. In the first version, we only varied *N_EE_* and *N_IE_* to dynamically modulate the circuit’s net E/I ratio (Fig 8B), in a way consistent with recent modeling of the effects of E/I ratio on a cortical circuit for perceptual decision-making [44]. In the second version, we varied *N_EE_*, *N_IE_*, and *N_EI_* (Fig S8A). Here, *N_EI_* was modulated independently from *N_EE_*, and *N_IE_*, which in turn were co-modulated by the same factor.

Per parameter combination, we ran 10 simulations, using the Brian2 spiking neural networks simulator [119]. Each simulation was run for 1000 seconds, with a random initialization of the network structure and the probabilistic spiking.

## ACKNOWLEDGEMENTS

The authors thank Christiane Reissmann for help with the data collection, as well as Sander Nieuwenhuis and Peter Murphy for helpful comments on the manuscript. This work was supported by the German Research Foundation (DFG): Heisenberg Professorship DO 1240/3-1 (to T.H.D.), and the Collaborative Research Center SFB 936 (Projects A2/A3, A7, Z3, to A.K.E., T.H.D., G.N., respectively); by the BMBF (Project 161A130, to A.K.E.); by the EU (ERC-2010-AdG-269716, to A.K.E.), by a grant from the state of Hamburg (CROSS FV25, to A. K.E.); and by the Netherlands Organization for Scientific Research (NWO, dossiernummer 406-15-256 to K.L.-H. and A.-E.A.)

## AUTHOR CONTRIBUTIONS

Conceptualization: T.P., A.K.E., and T.H.D.; Experimental design: T.P. and T.H.D.; Model design: T.P., A-E.A., K.L-H., and T.H.D.; Investigation: T.P.; Formal analysis: T.P.; Model simulations: A.-E.A.; Writing - Original draft: T.P. and T.H.D.; Writing – Review & Editing: T.P., A-E.A., G.N., A.K.E., K.L-H., and T.H.D. - Funding Acquisition: K.L-H., A.K.E., and T.H.D.; Supervision: G.N., K.L-H., and T.H.D.

## COMPTETING FINANCIAL INTERESTS

The authors declare no competing financial interests.

---

**S1 Fig.**
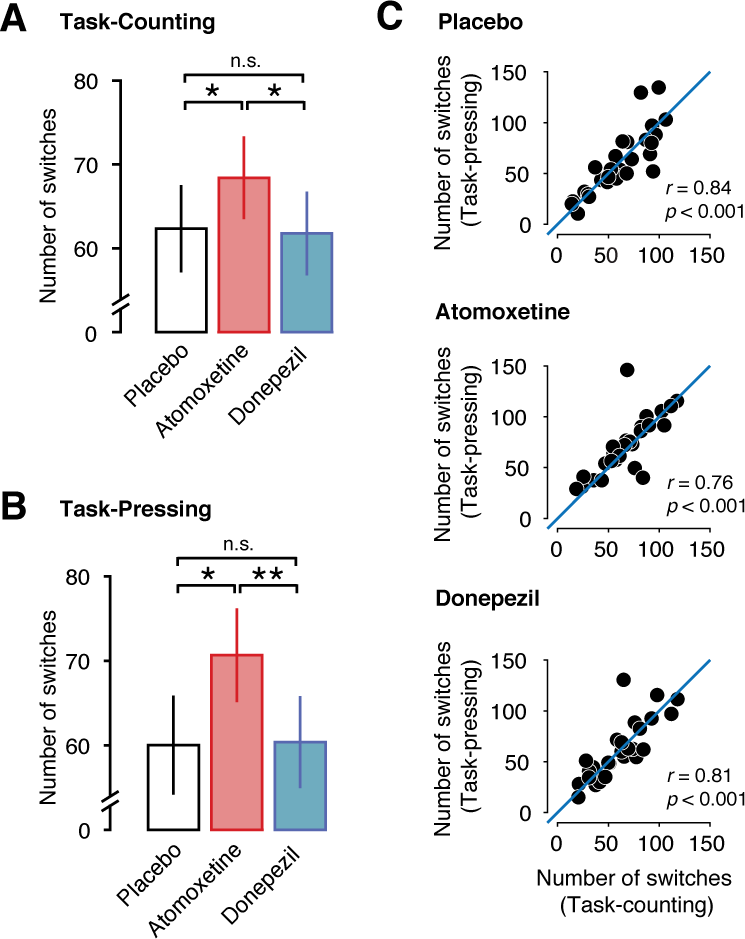
Similar Atomoxetine-related effects in both Task-counting and Task-resting conditions. **(A)** Number of perceptual alternations reported by the subjects per 10 min run for Task-counting condition. **(B)** Same as (A), but for Task-pressing condition. **(C)** Relation between the number of reported alternations during Task-counting (x-axis) and Task-pressing (y-axis). The blue line depicts a linear relation with slope 1 as a reference. Two-sided t-tests and Pearson correlations (N=28).

**S2 Fig.**
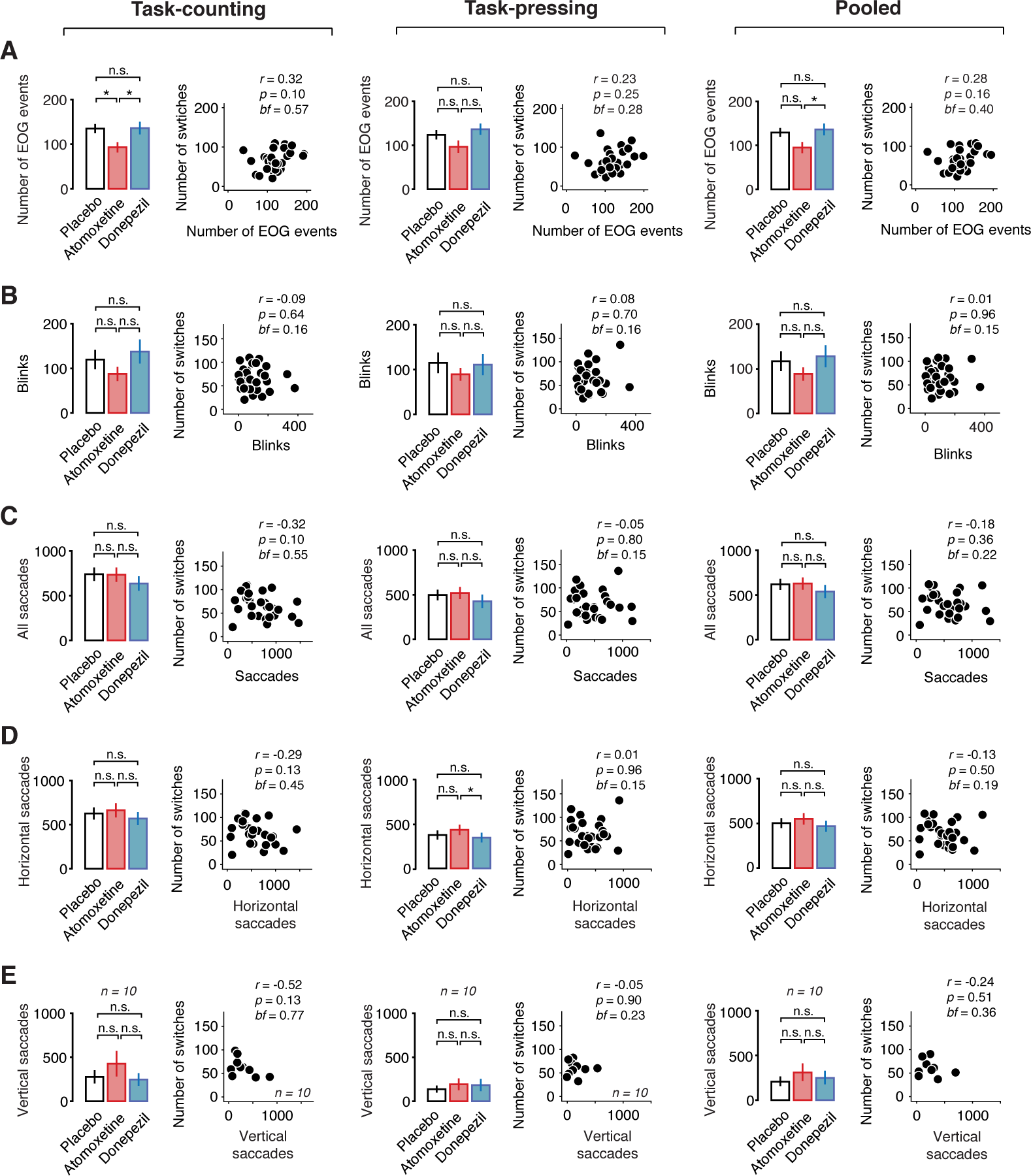
Change in perceptual alternation rate is not due to change in blinks or fixational eye movements. **(A)** Number of EOG events for during Task-counting (left), Taskpressing (middle) and pooled across both conditions (right). Scatter plots depict the relation between the number of EOG events (x-axis) and the number of reported perceptual alternations (y-axis). **(B)** Same as (A), but for the number of detected eye blinks. **(C)** Same as (A) and (B), but for the number of saccades (horizontal and vertical). **(D)** Same as (C), but for horizontal saccades only. **(E)** Same as (D), but for vertical saccades only. Two-sided t-tests and Pearson correlations (N=28). BF, Bayes factor. These control analyses demonstrate that the change in perceptual dynamics under Atomoxetine is not explained by changes in ocular parameters.

**S3 Fig.**
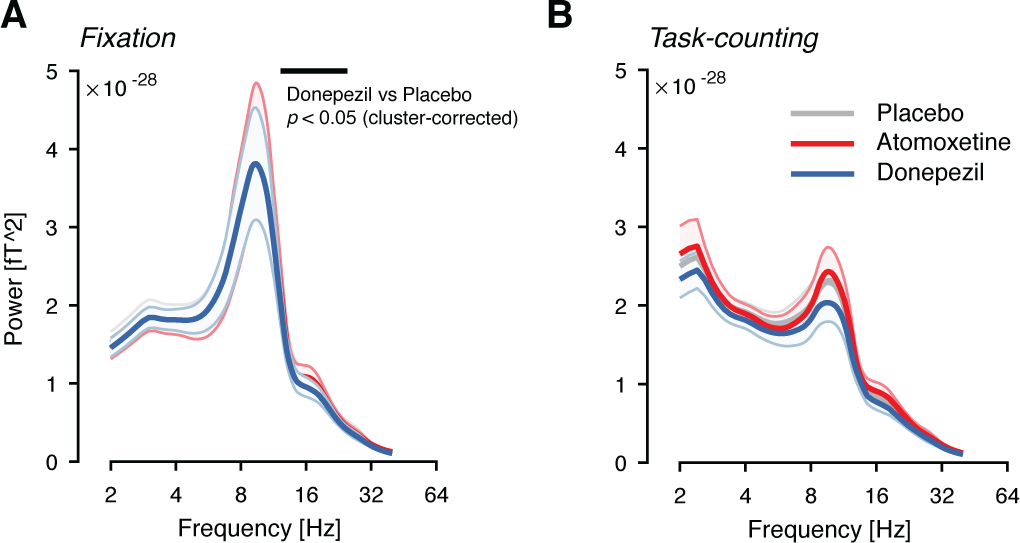
Power spectra, averaged across all MEG sensors, during Fixation **(A)** and Task-counting. **(B)** Black bar denotes significant differences assessed using a paired cluster-based permutation test (p < 0.05).

**S4 Fig.**
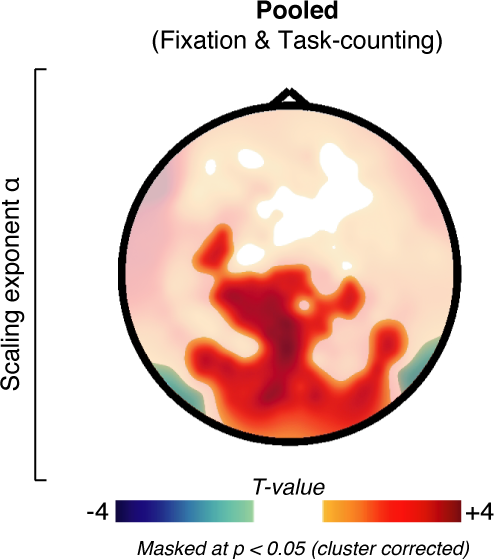
Sensor-level scaling exponent for the Atomoxetine condition, pooled across Fixation and Task-counting conditions. Thresholded at *p* = 0.05, two-sided cluster-based permutation test.

**S5 Fig.**
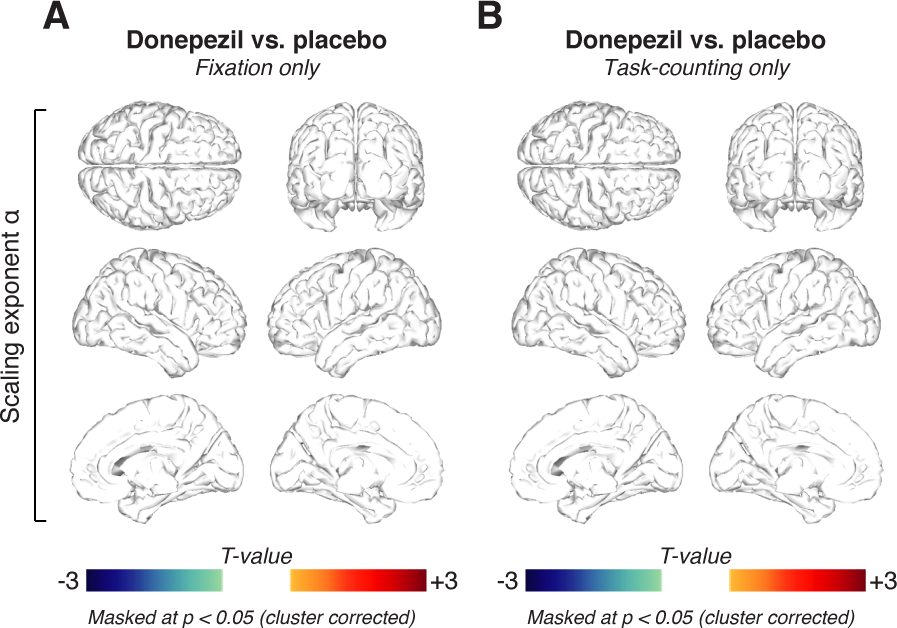
No Donepezil-related changes in scaling exponent in neither behavioral contexts.**(A)** Spatial distribution of Donepezil-induced changes in scaling exponent **α** during Fixation, thresholded at *p* = 0.05 (two-sided cluster-based permutation test). **(B)** As (A), but for Task-counting.

**S6 Fig.**
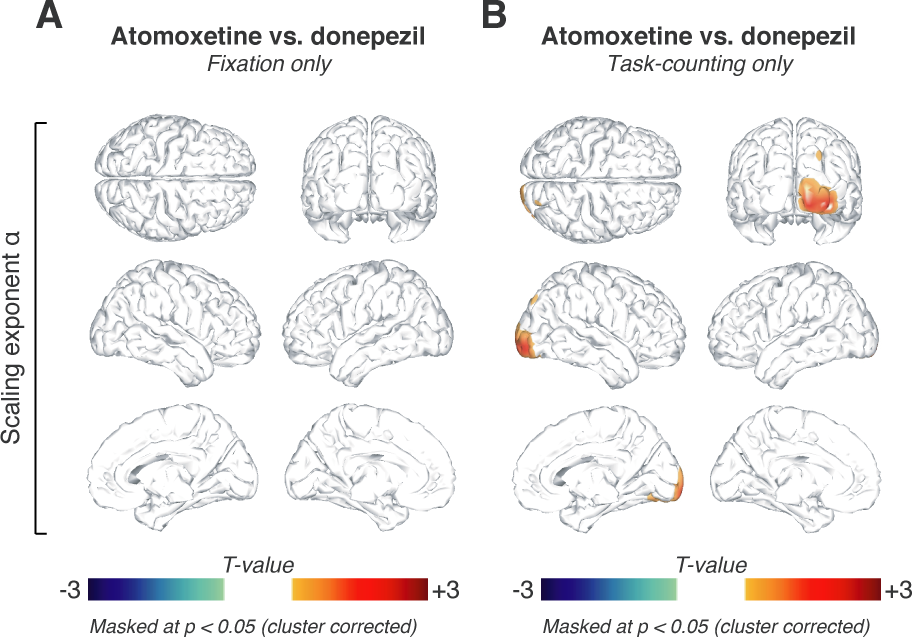
Direct comparison of the drug effects on scaling exponent **α**. **(A)** Comparison of the effects of the two drugs conditions (i.e., Atomoxetine vs. Donepezil) during Fixation.**(B)** Same as (A), but during Task-counting. All thresholds at *p* = 0.05, cluster-based two-sided permutation tests (N=28).

**S7 Fig.**
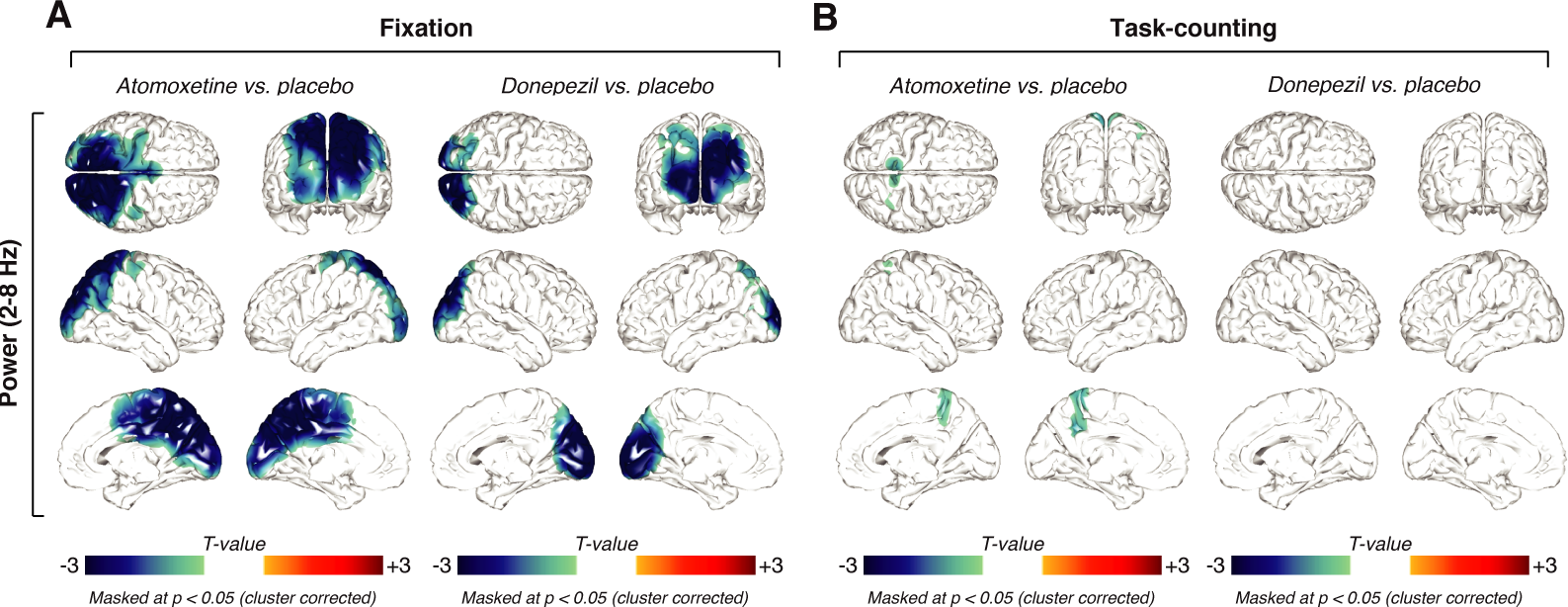
Similar effects of Atomoxetine and Donepezil on low-frequency (2-8 Hz) power. **(A)** Spatial distribution of drug-related low-frequency power changes during Fixation, thresholded at *p* = 0.05 (two-sided cluster-based permutation test). *Left*. Power changes after the administration of atomoxetine. *Right*. Power changes after the administration of donepezil. **(B)** Same as (A), but for Task-counting. The changes in low-frequency power, in combination with the reported decreases in alpha-band power, demonstrate a robust effect of both drugs on cortical dynamics.

**S8 Fig.**
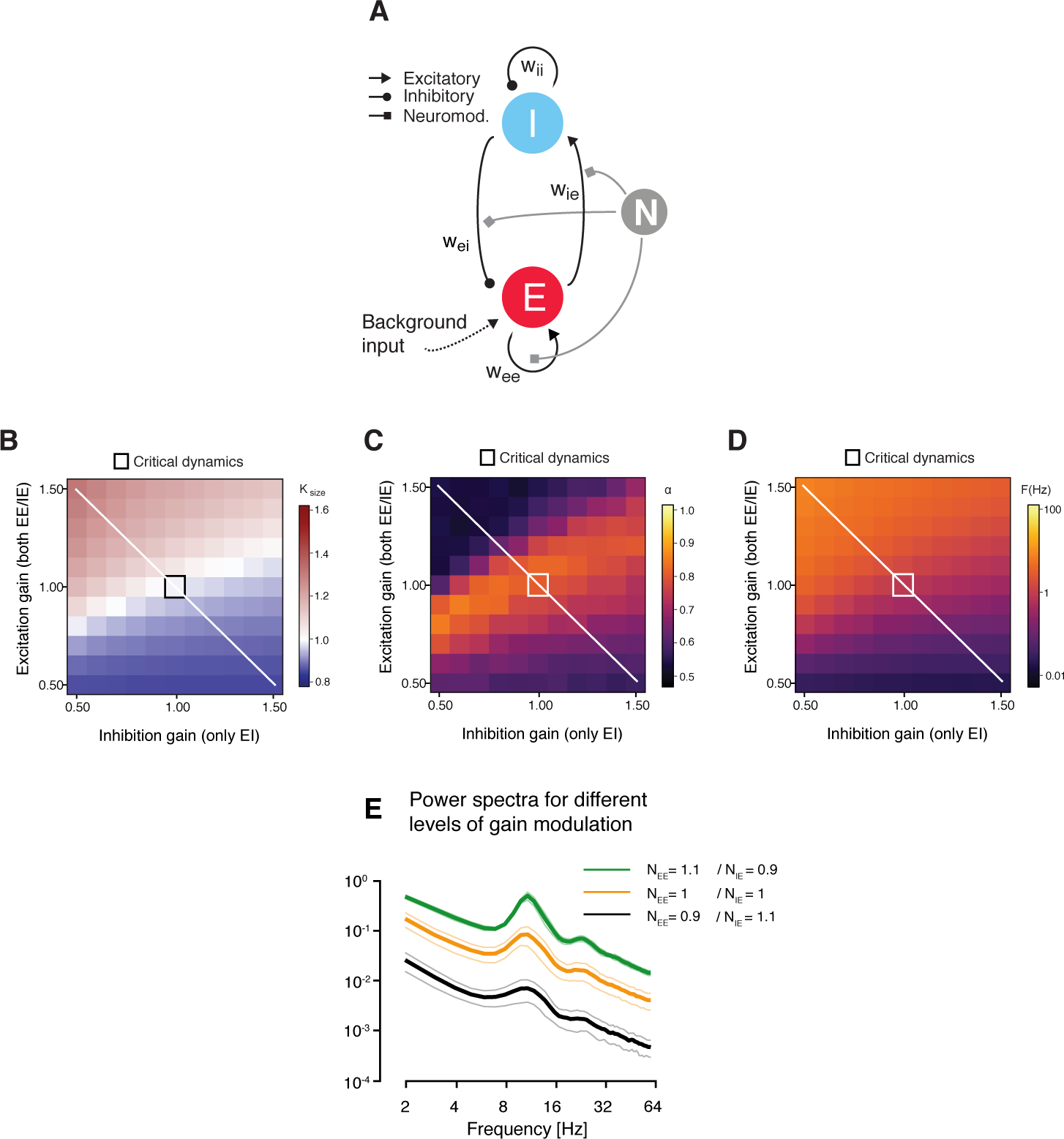
Different version of modulation of E/I ratio in cortical patch model **(A)** Neuromodulation was simulated as a gain modulation term multiplied with excitatory (EE and IE) and/or inhibitory (EI only) synaptic weights. **(B)** *κ* as a function of excitatory and inhibitory connectivity (with a spacing of 2.5%; means across 10 simulations per cell). The region of *κ*~1, overlaps with the region of *α* > 0.5 and splits the phase space into an excitation-dominant (k>1) and an inhibition-dominant region (*κ*<1). **(C)** Same as (B), but for scaling exponent *α*. **(D)** Same as (B) and (C), but for firing rate. In sum, the alternative version of modulation of E/I ratio yields comparable results to the version presented in Figure 8. **(E)** Model power spectra under different levels of synaptic gain modulation (neuromodulation).

## REFERENCES

1. Harris KD, Thiele A. Cortical state and attention. Nat Rev Neurosci. 2011;12: 509–523. doi:10.1038/nrn3084

2. McGinley MJ, Vinck M, Reimer J, Batista-Brito R, Zagha E, Cadwell CR, et al. Waking State: Rapid Variations Modulate Neural and Behavioral Responses. Neuron. 2015;87: 1143–1161. doi:10.1016/j.neuron.2015.09.012

3. Guedj C, Monfardini E, Reynaud AJ, Farnè A, Meunier M, Hadj-Bouziane F. Boosting Norepinephrine Transmission Triggers Flexible Reconfiguration of Brain Networks at Rest. Cereb Cortex. 2016; doi:10.1093/cercor/bhw262

4. van den Brink RL, Pfeffer T, Warren CM, Murphy PR, Tona K-D, van der Wee NJA, et al. Catecholaminergic Neuromodulation Shapes Intrinsic MRI Functional Connectivity in the Human Brain. J Neurosci. 2016;36: 7865–7876. doi:10.1523/JNEUROSCI.0744-16.2016

5. Aston-Jones G, Cohen JD. An integrative theory of locus coeruleus-norepinephrine function: adaptive gain and optimal performance. Annu Rev Neurosci. 2005;28: 403–450. doi:10.1146/annurev.neuro.28.061604.135709

6. Yu AJ, Dayan P. Uncertainty, neuromodulation, and attention. Neuron. 2005;46: 681–692. doi:10.1016/j.neuron.2005.04.026

7. Nelson A, Mooney R. The Basal Forebrain and Motor Cortex Provide Convergent yet Distinct Movement-Related Inputs to the Auditory Cortex. Neuron. 2016;90: 635–48. doi:10.1016/j.neuron.2016.03.031

8. de Gee JW, Colizoli O, Kloosterman NA, Knapen T, Nieuwenhuis S, Donner TH. Dynamic modulation of decision biases by brainstem arousal systems. eLife. 2017;6. doi:10.7554/eLife.23232

9. Berridge CW. Noradrenergic modulation of arousal. Brain Res Rev. 2008;58: 1–17. doi:10.1016/j.brainresrev.2007.10.013

10. Robbins TW, Arnsten AFT. The neuropsychopharmacology of fronto-executive function: monoaminergic modulation. Annu Rev Neurosci. 2009;32: 267–287. doi:10.1146/annurev.neuro.051508.135535

11. Lee S-H, Dan Y. Neuromodulation of brain states. Neuron. 2012;76: 209222.

12. Froemke RC. Plasticity of Cortical Excitatory-Inhibitory Balance. Annu Rev Neurosci. 2015;38: 195–219. doi:10.1146/annurev-neuro-071714-034002

13. Glimcher PW. Indeterminacy in brain and behavior. Annu Rev Psychol. 2005;56: 25–56. doi:10.1146/annurev.psych.55.090902.141429

14. Renart A, Machens CK. Variability in neural activity and behavior. Curr Opin Neurobiol. 2014;25: 211–220. doi:10.1016/j.conb.2014.02.013

15. Frank MJ, Doll BB, Oas-Terpstra J, Moreno F. Prefrontal and striatal dopaminergic genes predict individual differences in exploration and exploitation. Nat Neurosci. 2009;12: 1062–1068. doi:10.1038/nn.2342

16. Moreno-Bote R, Knill DC, Pouget A. Bayesian sampling in visual perception. Proc Natl Acad Sci. 2011;108: 12491–12496. doi:10.1073/pnas.1101430108

17. Pinto L, Goard MJ, Estandian D, Xu M, Kwan AC, Lee S-HH, et al. Fast modulation of visual perception by basal forebrain cholinergic neurons. 2013;16: 1857–63. doi:10.1038/nn.3552

18. Chen N, Sugihara H, Sur M. An acetylcholine-activated microcircuit drives temporal dynamics of cortical activity. Nat Neurosci. 2015;18: 892–902. doi:10.1038/nn.4002

19. Minces V, Pinto L, Dan Y, Chiba AA. Cholinergic shaping of neural correlations. Proc Natl Acad Sci. 2017;114: 5725–5730. doi:10.1073/pnas.1621493114

20. Polack P-O, Friedman J, Golshani P. Cellular mechanisms of brain state-dependent gain modulation in visual cortex. Nat Neurosci. 2013;16: 1331–1339. doi:10.1038/nn.3464

21. Fu Y, Tucciarone JM, Espinosa JS, Sheng N, Darcy DP, Nicoll RA, et al. A cortical circuit for gain control by behavioral state. Cell. 2014;156: 1139–1152. doi:10.1016/j.cell.2014.01.050

22. Fox MD, Raichle ME. Spontaneous fluctuations in brain activity observed with functional magnetic resonance imaging. Nat Rev Neurosci. 2007;8: 700–711. doi:10.1038/nrn2201

23. Deco G, Jirsa VK, McIntosh AR. Emerging concepts for the dynamical organization of resting-state activity in the brain. Nat Rev Neurosci. 2011;12: 43–56. doi:10.1038/nrn2961

24. Miller KJ, Sorensen LB, Ojemann JG, den Nijs M. Power-law scaling in the brain surface electric potential. PLoS Comput Biol. 2009;5: e1000609. doi:10.1371/journal.pcbi.1000609

25. He BJ, Zempel JM, Snyder AZ, Raichle ME. The temporal structures and functional significance of scale-free brain activity. Neuron. 2010;66: 353–369. doi:10.1016/j.neuron.2010.04.020

26. Linkenkaer-Hansen K, Nikouline VV, Palva JM, Ilmoniemi RJ. Long-range temporal correlations and scaling behavior in human brain oscillations. J Neurosci Off J Soc Neurosci. 2001;21: 1370–1377.

27. He BJ. Scale-Free Properties of the Functional Magnetic Resonance Imaging Signal during Rest and Task. J Neurosci. 2011;31: 13786–13795. doi:10.1523/JNEUROSCI.2111-11.2011

28. Palva JM, Zhigalov A, Hirvonen J, Korhonen O, Linkenkaer-Hansen K, Palva S. Neuronal long-range temporal correlations and avalanche dynamics are correlated with behavioral scaling laws. Proc Natl Acad Sci. 2013;110: 3585–3590. doi:10.1073/pnas.1216855110

29. Zhigalov A, Arnulfo G, Nobili L, Palva S, Palva JM. Relationship of fast-and slow-timescale neuronal dynamics in human MEG and SEEG. J Neurosci Off J Soc Neurosci. 2015;35: 5385–5396. doi:10.1523/JNEUROSCI.4880-14.2015

30. Honey CJ, Thesen T, Donner TH, Silbert LJ, Carlson CE, Devinsky O, et al. Slow Cortical Dynamics and the Accumulation of Information over Long Timescales. Neuron. 2012;76: 423–434. doi:10.1016/j.neuron.2012.08.011

31. Donner TH, Sagi D, Bonneh YS, Heeger DJ. Retinotopic Patterns of Correlated Fluctuations in Visual Cortex Reflect the Dynamics of Spontaneous Perceptual Suppression. J Neurosci. 2013;33: 2188–2198. doi:10.1523/JNEUROSCI.3388-12.2013

32. Chaudhuri R, Knoblauch K, Gariel M-A, Kennedy H, Wang X-J. A Large-Scale Circuit Mechanism for Hierarchical Dynamical Processing in the Primate Cortex. Neuron. 2015;88: 419–431. doi:10.1016/j.neuron.2015.09.008

33. Moreno-Bote R, Rinzel J, Rubin N. Noise-induced alternations in an attractor network model of perceptual bistability. J Neurophysiol. 2007;98: 1125–1139. doi:10.1152/jn.00116.2007

34. Noest AJ, van Ee R, Nijs MM, van Wezel RJA. Percept-choice sequences driven by interrupted ambiguous stimuli: A low-level neural model. J Vis. 2007;7: 10. doi:10.1167/7.8.10

35. Deco G, Romo R. The role of fluctuations in perception. Trends Neurosci. 2008;31: 591–598. doi:10.1016/j.tins.2008.08.007

36. van Loon AM, Knapen T, Scholte HS, St. John-Saaltink E, Donner TH, Lamme VAF. GABA Shapes the Dynamics of Bistable Perception. Curr Biol. 2013;23: 823–827. doi:10.1016/j.cub.2013.03.067

37. Eckhoff P, Wong-Lin KF, Holmes P. Optimality and Robustness of a Biophysical Decision-Making Model under Norepinephrine Modulation. J Neurosci. 2009;29: 4301–4311. doi:10.1523/JNEUROSCI.5024-08.2009

38. Parker AJ, Krug K, Cumming BG. Neuronal activity and its links with the perception of multi-stable figures. Philos Trans R Soc Lond B Biol Sci. 2002;357: 1053–1062. doi:10.1098/rstb.2002.1112

39. Brouwer GJ, van Ee R. Visual cortex allows prediction of perceptual states during ambiguous structure-from-motion. J Neurosci Off J Soc Neurosci. 2007;27: 1015–1023. doi:10.1523/JNEUROSCI.4593-06.2007

40. Donner TH, Siegel M. A framework for local cortical oscillation patterns. Trends Cogn Sci. 2011;15: 191–199. doi:10.1016/j.tics.2011.03.007

41. Linkenkaer-Hansen K, Smit DJA, Barkil A, van Beijsterveldt TEM, Brussaard AB, Boomsma DI, et al. Genetic contributions to long-range temporal correlations in ongoing oscillations. J Neurosci Off J Soc Neurosci. 2007;27: 13882–13889. doi:10.1523/JNEUROSCI.3083-07.2007

42. Linkenkaer-Hansen K, Nikulin VV, Palva S, Ilmoniemi RJ, Palva JM. Prestimulus oscillations enhance psychophysical performance in humans. J Neurosci Off J Soc Neurosci. 2004;24: 10186–10190. doi:10.1523/JNEUROSCI.2584-04.2004

43. Bauer M, Kluge C, Bach D, Bradbury D, Heinze HJ, Dolan RJ, et al. Cholinergic enhancement of visual attention and neural oscillations in the human brain. Curr Biol CB. 2012;22: 397–402. doi:10.1016/j.cub.2012.01.022

44. Lam NH, Borduqui T, Hallak J, Roque AC, Anticevic A, Krystal JH, et al. Effects of Altered Excitation-Inhibition Balance on Decision Making in a Cortical Circuit Model. bioRxiv. 2017; doi:http://dx.doi.org/10.1101/100347

45. van Vreeswijk C, Sompolinsky H. Chaos in neuronal networks with balanced excitatory and inhibitory activity. Science. 1996;274: 1724–1726.

46. Shadlen MN, Newsome WT. The variable discharge of cortical neurons: implications for connectivity, computation, and information coding. J Neurosci Off J Soc Neurosci. 1998;18: 3870–3896.

47. Okun M, Lampl I. Balance of excitation and inhibition. Scholarpedia. 2009;4: 7467. doi:10.4249/scholarpedia.7467

48. Isaacson JS, Scanziani M. How inhibition shapes cortical activity. Neuron. 2011;72: 231–243. doi:10.1016/j.neuron.2011.09.027

49. Poil S-S, Hardstone R, Mansvelder HD, Linkenkaer-Hansen K. Critical-State Dynamics of Avalanches and Oscillations Jointly Emerge from Balanced Excitation/Inhibition in Neuronal Networks. J Neurosci. 2012;32: 9817–9823. doi:10.1523/JNEUROSCI.5990-11.2012

50. Beggs JM, Plenz D. Neuronal avalanches in neocortical circuits. J Neurosci Off J Soc Neurosci. 2003;23: 11167–11177.

51. Servan-Schreiber D, Printz H, Cohen J. A network model of catecholamine effects: gain, signal-to-noise ratio, and behavior. Science. 1990;249: 892–895. doi:10.1126/science.2392679

52. Haider B, Häusser M, Carandini M. Inhibition dominates sensory responses in the awake cortex. Nature. 2013;493: 97–100. doi:10.1038/nature11665

53. Adesnik H. Synaptic Mechanisms of Feature Coding in the Visual Cortex of Awake Mice. Neuron. 2017;95: 1147–1159.e4. doi:10.1016/j.neuron.2017.08.014

54. Sara SJ. The locus coeruleus and noradrenergic modulation of cognition. Nat Rev Neurosci. 2009;10: 211–223. doi:10.1038/nrn2573

55. Morrison JH, Foote SL. Noradrenergic and serotoninergic innervation of cortical, thalamic, and tectal visual structures in Old and New World monkeys. J Comp Neurol. 1986;243: 117–138. doi:10.1002/cne.902430110

56. Pennartz CM. The ascending neuromodulatory systems in learning by reinforcement: comparing computational conjectures with experimental findings. Brain Res Brain Res Rev. 1995;21: 219–245.

57. Roelfsema PR, van Ooyen A, Watanabe T. Perceptual learning rules based on reinforcers and attention. Trends Cogn Sci. 2010;14: 64–71. doi:10.1016/j.tics.2009.11.005

58. Salgado H, Treviño M, Atzori M. Layer- and area-specific actions of norepinephrine on cortical synaptic transmission. Brain Res. 2016;1641: 163–176. doi:10.1016/j.brainres.2016.01.033

59. Ramos BP, Arnsten AFT. Adrenergic pharmacology and cognition: focus on the prefrontal cortex. Pharmacol Ther. 2007;113: 523–536. doi:10.1016/j.pharmthera.2006.11.006

60. Crochet S, Poulet JFA, Kremer Y, Petersen CCH. Synaptic Mechanisms Underlying Sparse Coding of Active Touch. Neuron. 2011;69: 1160–1175. doi:10.1016/j.neuron.2011.02.022

61. Zhou M, Liang F, Xiong XR, Li L, Li H, Xiao Z, et al. Scaling down of balanced excitation and inhibition by active behavioral states in auditory cortex. Nat Neurosci. 2014;17: 841–850. doi:10.1038/nn.3701

62. Swadlow HA. Thalamocortical control of feed-forward inhibition in awake somatosensory “barrel” cortex. Philos Trans R Soc B Biol Sci. 2002;357: 1717–1727. doi:10.1098/rstb.2002.1156

63. Kepecs A, Fishell G. Interneuron cell types are fit to function. Nature. 2014;505: 318–326. doi:10.1038/nature12983

64. Ponce-Alvarez A, He BJ, Hagmann P, Deco G. Task-Driven Activity Reduces the Cortical Activity Space of the Brain: Experiment and Whole-Brain Modeling. Graham LJ, editor. PLOS Comput Biol. 2015;11: e1004445. doi:10.1371/journal.pcbi.1004445

65. Froemke RC, Merzenich MM, Schreiner CE. A synaptic memory trace for cortical receptive field plasticity. Nature. 2007;450: 425–429. doi:10.1038/nature06289

66. Martins ARO, Froemke RC. Coordinated forms of noradrenergic plasticity in the locus coeruleus and primary auditory cortex. Nat Neurosci. 2015;18: 1483–1492. doi:10.1038/nn.4090

67. Murphy BK, Miller KD. Multiplicative gain changes are induced by excitation or inhibition alone. J Neurosci. 2003;23: 10040–10051.

68. Denève S, Machens CK. Efficient codes and balanced networks. Nat Neurosci. 2016;19: 375–382. doi:10.1038/nn.4243

69. Wang X-J. Decision making in recurrent neuronal circuits. Neuron. 2008;60: 215–234. doi:10.1016/j.neuron.2008.09.034

70. Yizhar O, Fenno LE, Prigge M, Schneider F, Davidson TJ, O’Shea DJ, et al. Neocortical excitation/inhibition balance in information processing and social dysfunction. Nature. 2011;477: 171–178. doi:10.1038/nature10360

71. Lisman J. Excitation, inhibition, local oscillations, or large-scale loops: what causes the symptoms of schizophrenia? Curr Opin Neurobiol. 2012;22: 537–544. doi:10.1016/j.conb.2011.10.018

72. Nelson SB, Valakh V. Excitatory/Inhibitory Balance and Circuit Homeostasis in Autism Spectrum Disorders. Neuron. 2015;87: 684–698. doi:10.1016/j.neuron.2015.07.033

73. Fuchs T, Jefferson SJ, Hooper A, Yee P-H, Maguire J, Luscher B. Disinhibition of somatostatin-positive GABAergic interneurons results in an anxiolytic and antidepressant-like brain state. Mol Psychiatry. 2017;22: 920–930. doi:10.1038/mp.2016.188

74. Eggermann E, Kremer Y, Crochet S, Petersen CCH. Cholinergic Signals in Mouse Barrel Cortex during Active Whisker Sensing. Cell Rep. 2014;9: 1654–1660. doi:10.1016/j.celrep.2014.11.005

75. Carter OL, Pettigrew JD, Hasler F, Wallis GM, Liu GB, Hell D, et al. Modulating the rate and rhythmicity of perceptual rivalry alternations with the mixed 5-HT2A and 5-HT1A agonist psilocybin. Neuropsychopharmacol Off Publ Am Coll Neuropsychopharmacol. 2005;30: 1154–1162. doi:10.1038/sj.npp.1300621

76. Noudoost B, Moore T. Control of visual cortical signals by prefrontal dopamine. Nature. 2011;474: 372–375. doi:10.1038/nature09995

77. Murray JD, Bernacchia A, Freedman DJ, Romo R, Wallis JD, Cai X, et al. A hierarchy of intrinsic timescales across primate cortex. Nat Neurosci. 2014;17: 1661–1663. doi:10.1038/nn.3862

78. Wang X-J. Synaptic reverberation underlying mnemonic persistent activity. 2001;24. doi:10.1016/s0166-2236(00)01868-3

79. Wang X-JJ. Probabilistic decision making by slow reverberation in cortical circuits. 2002;36: 955–68. doi:10.1016/s0896-6273(02)01092-9

80. Beggs JM. The criticality hypothesis: how local cortical networks might optimize information processing. Philos Transact A Math Phys Eng Sci. 2008;366: 329–343. doi:10.1098/rsta.2007.2092

81. Bak P, Tang C, Wiesenfeld K. Self-organized criticality: An explanation of the 1/f noise. Phys Rev Lett. 1987;59: 381–384. doi:10.1103/PhysRevLett.59.381

82. Bak P. How nature works: the science of self-organized criticality [Internet]. New York, NY, USA: Copernicus; 1996. Available: http://catalog.hathitrust.org/api/volumes/oclc/34623628.html

83. Chialvo DR. Emergent complex neural dynamics. Nat Phys. 2010;6: 744–750. doi:10.1038/nphys1803

84. Hesse J, Gross T. Self-organized criticality as a fundamental property of neural systems. Front Syst Neurosci. 2014;8: 166. doi:10.3389/fnsys.2014.00166

85. Kinouchi O, Copelli M. Optimal dynamical range of excitable networks at criticality. Nat Phys. 2006;2: 348–351. doi:10.1038/nphys289

86. Shew WL, Yang H, Petermann T, Roy R, Plenz D. Neuronal avalanches imply maximum dynamic range in cortical networks at criticality. J Neurosci Off J Soc Neurosci. 2009;29: 15595–15600. doi:10.1523/JNEUROSCI.3864-09.2009

87. Shew W, Yang H, Yu S, Roy R. Information capacity and transmission are maximized in balanced cortical networks with neuronal avalanches. J …. 2011;

88. Shriki O, Yellin D. Optimal Information Representation and Criticality in an Adaptive Sensory Recurrent Neuronal Network. PLoS Comput Biol. 2016;12: e1004698. doi:10.1371/journal.pcbi.1004698

89. Priesemann V, Valderrama M, Wibral M, Le Van Quyen M. Neuronal avalanches differ from wakefulness to deep sleep--evidence from intracranial depth recordings in humans. PLoS Comput Biol. 2013;9: e1002985. doi:10.1371/journal.pcbi.1002985

90. Arviv O, Goldstein A, Shriki O. Near-Critical Dynamics in Stimulus-Evoked Activity of the Human Brain and Its Relation to Spontaneous Resting-State Activity. J Neurosci Off J Soc Neurosci. 2015;35: 13927–13942. doi:10.1523/JNEUROSCI.0477-15.2015

91. Fagerholm ED, Lorenz R, Scott G, Dinov M, Hellyer PJ, Mirzaei N, et al. Cascades and Cognitive State: Focused Attention Incurs Subcritical Dynamics. J Neurosci. 2015;35: 4626–4634. doi:10.1523/JNEUROSCI.3694-14.2015

92. Shew WL, Clawson WP, Pobst J, Karimipanah Y, Wright NC, Wessel R. Adaptation to sensory input tunes visual cortex to criticality. Nat Phys. 2015;11: 659–663. doi:10.1038/nphys3370

93. Krug K, Cicmil N, Parker AJ, Cumming BG. A Causal Role for V5/MT Neurons Coding Motion-Disparity Conjunctions in Resolving Perceptual Ambiguity. 2013;23: 1454–9. doi:10.1016/j.cub.2013.06.023

94. Bollimunta A, Mo J, Schroeder CE, Ding M. Neuronal mechanisms and attentional modulation of corticothalamic α oscillations. J Neurosci Off J Soc Neurosci. 2011;31: 4935–4943. doi:10.1523/JNEUROSCI.5580-10.2011

95. Bymaster FP, Katner JS, Nelson DL, Hemrick-Luecke SK, Threlkeld PG, Heiligenstein JH, et al. Atomoxetine increases extracellular levels of norepinephrine and dopamine in prefrontal cortex of rat: a potential mechanism for efficacy in attention deficit/hyperactivity disorder. Neuropsychopharmacol Off Publ Am Coll Neuropsychopharmacol. 2002;27: 699–711. doi:10.1016/S0893-133X(02)00346-9

96. Ding Y-S, Naganawa M, Gallezot J-D, Nabulsi N, Lin S-F, Ropchan J, et al. Clinical doses of atomoxetine significantly occupy both norepinephrine and serotonin transports: Implications on treatment of depression and ADHD. NeuroImage. 2014;86: 164–171. doi:10.1016/j.neuroimage.2013.08.001

97. Tiseo, Rogers, Friedhoff. Pharmacokinetic and pharmacodynamic profile of donepezil HCl following evening administration: Evening administration of donepezil HCl. Br J Clin Pharmacol. 1998;46: 13–18. doi:10.1046/j.1365-2125.1998.0460s1013.x

98. Sauer J-M, Ring BJ, Witcher JW. Clinical pharmacokinetics of atomoxetine. Clin Pharmacokinet. 2005;44: 571–590. doi:10.2165/00003088-200544060-00002

99. Wallach H, O’connell DN. The kinetic depth effect. J Exp Psychol. 1953;45: 205–217.

100. Sperling G, Dosher BA, Landy MS. How to study the kinetic depth effect experimentally. J Exp Psychol Hum Percept Perform. 1990;16: 445–450.

101. Engbert R, Kliegl R. Microsaccades uncover the orientation of covert attention. Vision Res. 2003;43: 1035–1045.

102. Oostenveld R, Fries P, Maris E, Schoffelen J-M. FieldTrip: Open source software for advanced analysis of MEG, EEG, and invasive electrophysiological data. Comput Intell Neurosci. 2011;2011: 156869. doi:10.1155/2011/156869

103. Bell AJ, Sejnowski TJ. An information-maximization approach to blind separation and blind deconvolution. Neural Comput. 1995;7: 1129–1159.

104. Hyvarinen A. Fast and robust fixed-point algorithms for independent component analysis. IEEE Trans Neural Netw. 1999;10: 626–634. doi:10.1109/72.761722

105. Hipp JF, Siegel M. Dissociating neuronal gamma-band activity from cranial and ocular muscle activity in EEG. Front Hum Neurosci. 2013;7. doi:10.3389/fnhum.2013.00338

106. Mitra PP, Pesaran B. Analysis of dynamic brain imaging data. Biophys J. 1999;76: 691–708. doi:10.1016/S0006-3495(99)77236-X

107. Pascual-Marqui RD, Lehmann D, Koukkou M, Kochi K, Anderer P, Saletu B, et al. Assessing interactions in the brain with exact low-resolution electromagnetic tomography. Philos Transact A Math Phys Eng Sci. 2011;369: 3768–3784. doi:10.1098/rsta.2011.0081

108. Fonov V, Evans AC, Botteron K, Almli CR, McKinstry RC, Collins DL, et al. Unbiased average age-appropriate atlases for pediatric studies. NeuroImage. 2011;54: 313–327. doi:10.1016/j.neuroimage.2010.07.033

109. Nolte G. The magnetic lead field theorem in the quasi-static approximation and its use for magnetoencephalography forward calculation in realistic volume conductors. Phys Med Biol. 2003;48: 3637–3652.

110. Pfeffer T, Linkenkaer-Hansen K, Avramiea A-E, Engel AK, Donner TH. Noradrenaline increases long-range temporal correlations of neuronal alpha oscillations in the human cortex. 2015. p. 393.27.

111. Peng CK, Buldyrev SV, Havlin S, Simons M, Stanley HE, Goldberger AL. Mosaic organization of DNA nucleotides. Phys Rev E Stat Phys Plasmas Fluids Relat Interdiscip Top. 1994;49: 1685–1689.

112. Hardstone R, Poil S-S, Schiavone G, Jansen R, Nikulin VV, Mansvelder HD, et al. Detrended Fluctuation Analysis: A Scale-Free View on Neuronal Oscillations. Front Physiol. 2012;3. doi:10.3389/fphys.2012.00450

113. Montez T, Poil S-S, Jones BF, Manshanden I, Verbunt JPA, van Dijk BW, et al. Altered temporal correlations in parietal alpha and prefrontal theta oscillations in early-stage Alzheimer disease. Proc Natl Acad Sci. 2009;106: 1614–1619. doi:10.1073/pnas.0811699106

114. Rouder JN, Speckman PL, Sun D, Morey RD, Iverson G. Bayesian t tests for accepting and rejecting the null hypothesis. Psychon Bull Rev. 2009;16: 225–237. doi:10.3758/PBR.16.2.225

115. Wetzels R, Wagenmakers E-J. A default Bayesian hypothesis test for correlations and partial correlations. Psychon Bull Rev. 2012;19: 10571064. doi:10.3758/s13423-012-0295-x

116. Nichols TE, Holmes AP. Nonparametric permutation tests for functional neuroimaging: A primer with examples. Hum Brain Mapp. 2002;15: 1–25. doi:10.1002/hbm.1058

117. Maris E, Oostenveld R. Nonparametric statistical testing of EEG- and MEG-data. J Neurosci Methods. 2007;164: 177–190. doi:10.1016/j.jneumeth.2007.03.024

118. Eiben AE, Smit SK. Parameter tuning for configuring and analyzing evolutionary algorithms. Swarm Evol Comput. 2011;1: 19–31. doi:10.1016/j.swevo.2011.02.001

119. Goodman DF, Stimberg M, Yger P, Brette R. Brian 2: neural simulations on a variety of computational hardware. BMC Neurosci. 2014;15: P199. doi:10.1186/1471-2202-15-S1-P199

